# Exploring genetic, expression and regulatory patterns of parental alleles in Muscovy duck (*Cairina moschata*) using haplotype-resolved assemblies

**DOI:** 10.64898/2026.03.04.709678

**Authors:** Te Li, Yiming Wang, Zhou Zhang, Chao Chen, Nenzhu Zheng, Jun Wang, Mengfei Ning, Jian Wang, Huashui Ai, Yinhua Huang

## Abstract

**Background:** Although the biological mechanism for heterosis has been debated for a long time, heterosis is widely utilized to increase the global productivity of crops and livestock. Recently, the mechanism has been well characterized in crops and livestock with a male-heterogametic XY system due to genomic assembly advancements, especially the availability of haploid genomes. However, the biological mechanism for heterosis remains unclear in poultry possessing the female-heterogametic ZW system.

**Results:** Here, we assembled chromosome-level diploid and haploid genomes of the Muscovy duck. We developed an efficient and cost-effective method to assemble 12 variation graph-haploid Muscovy duck genomes from three full-sibling pairs with high quality using short-read Illumina sequences. We further characterized genetic, expression and regulatory patterns of parental alleles at multiple scales. We found that maternal haploid genomes generally had more open chromatin organization and higher accessibility, and higher levels of gene expression, while showing similar DNA methylation levels when compared to paternal haploid genomes. In contrast, the female paternal Z chromosome showed the most, and the male paternal Z chromosome presented more, relaxed chromatin organization and chromatin accessibility, and gene expression compared to the male maternal Z chromosome. Thus, the ZW system largely relies on compensation and balance to regulate gene expression on the sex Z chromosome. Moreover, we identified non-Mendelian regions covering 0.26% of the genome (∼3.18 Mb). These regions contained lower gene density, GC content, and repeat sequence frequency, but were enriched for DNA motifs bound by transcription factors, likely leading to a compacted chromatin structure and lower chromatin accessibility.

**Conclusions:** Our work here provides a comprehensive profile of parental alleles genetic, expression and regulatory patterns in the female-heterogametic ZW system, and might be useful for the utilization of heterosis in poultry.

## Background

Hybrid breeding has made remarkable achievements in crops and livestock by generating heterosis, which is economically important to agricultural production. After a century of extensive research, observation and debate, three classical hypotheses including dominance [1], overdominance [2, 3], and epistasis [4, 5] based on quantitative trait loci (QTL) mapping and genome-wide association studies (GWAS) have been proposed for the genetic basis of heterosis. Recently, significant progress has been made in the genetic basis of heterosis in the last two decades with the advancement of genomic technologies, especially the availability of telomere-to-telomere diploid and haploid genomes. Such progress enables the characterization of patterns of paternal and maternal alleles on a genome-wide scale at multiple layers, including expression, chromatin conformation, chromatin accessibility and DNA methylation. In crops, integrating genomics with epigenomics (DNA methylation) has allowed researchers to unravel how the parent-of-origin specific expression of key alleles regulate critical yield traits like grain number and filling rate [6]. In livestock, effect of parental allele expression patterns to complex production traits such as muscle development (meat yield) and feed efficiency had been estimated using multiple omics data [7, 8]. A classic example of such large-scale parental alleles regulation in females of male-heterogametic (XY) mammals is X-chromosome inactivation (XCI). This presented a crucial role in sex-specific differences of fertility and disease susceptibility [9].

Unlike livestock possessing a male-heterogametic XY system, poultry are characterized by a female-heterogametic ZW system. In this system, sex chromosome inactivation analogous to X-inactivation has not been observed. Instead, birds often lack complete dosage compensation, with only a subset of genes exhibiting similar expression levels between sexes [10, 11]. The chicken was found to utilize a Z-linked microRNA (miR-2954) with strong male-biased expression to degrade transcripts of two upregulated dosage-sensitive Z gene copies in males [12]. These observations suggested that the avian female-heterogametic ZW system might show different expression and regulatory patterns of parental alleles when compared to the mammalian male-heterogametic XY system [10, 13]. However, parental allele expression and regulatory patterns in the ZW system remain unclear, partially limited by the availability of chromosome-level haploid genomes.

In this study, we generated highly contiguous chromosome-level Muscovy duck diploid and haploid genomes using 110-fold Nanopore long reads, 100-fold 150 bp paired-end (PE) Illumina genomic reads, 120-fold optical map molecules, 250-fold PE150 Hi-C reads and 100-fold parental PE150 genomic reads. We built a new efficient and cost-effective pipeline to assemble 12 VG-haploid haploid genomes and developed a method to evaluate the genetic inheritance of paternal and maternal alleles on a genome-wide scale. We then investigated patterns of parental alleles at multiple scales including genetics, expression and regulation.

## Methods

### Sample collection and DNA extraction

For the genomic analysis, genomic DNA was extracted from muscle tissue and fresh blood of Muscovy duck individuals using the QIAGEN Genomic DNA Kit (QIAGEN, Hilden, Germany) following the manufacturer’s instructions.

### Genome sequencing, assembly, and evaluation

To construct a high-continuity genome, a multi-platform sequencing strategy was implemented. For the generation of primary contigs, long-read libraries with an insert size greater than 20 kb were sequenced on the Oxford Nanopore Technologies (ONT) GridION X5 platform, supplemented by ultra-long reads generated on the PromethION platform. Simultaneously, for hybrid scaffolding, high-molecular-weight DNA was labeled with DLE-1 (8–25 labels/100 kb) and sequenced on the Bionano Saphyr instrument, retaining only molecules longer than 150 kb. Complementing these long-read technologies, short-read sequencing was also performed on the Illumina HiSeq X Ten platform to support multiple analytical goals. Standard WGS libraries (400 bp insert) were sequenced for genome polishing, while Hi-C libraries were constructed using HindIII digestion and streptavidin enrichment to facilitate chromosomal anchoring. Furthermore, ATAC-seq libraries were generated via Tn5 transposition to profile chromatin accessibility.

To construct a high-continuity genome, genome size (1.27 Gb) and heterozygosity were estimated via k-mer analysis (k = 21) using Jellyfish (v2.3.0) [14] and GenomeScope 2.0 [15]. For primary assembly, raw Nanopore reads were filtered (length ≥ 5 kb, q-value ≥ 7) and assembled using NextDenovo (v2.5.0) [16], followed by contig polishing with Nextpolish (v1.4.0) [17] using Illumina reads. To elevate contiguity, hybrid scaffolding was achieved by integrating Bionano optical maps via a custom R-based pipeline (scripts available on Figshare) and the SOLVE RefAligner (v3.2.1), with further refinement performed in Bionano Access (v1.7) [18]. Finally, chromosome-level anchoring was subsequently completed using Hi-C data: reads were processed by Trimmatic (v0.39) [19], mapped using BWA [20] and Juicer (v1.5) [21], and scaffolds were anchored using 3D-DNA (v180922) [22]. Upon completion of automated scaffolding, contact maps were manually curated in Juicebox (v1.11.08) [23]. The final assembly underwent three rounds of gap filling using GapCloser (v1.12) [24] with both standard and ultra-long Nanopore reads.

Following assembly, genomic integrity was assessed by comparing completeness against existing avian assemblies (SKLA2.0, KizCaiMos1.0, GRCg7b, bTaeGut1_v1), mapping RNA-seq reads using HISAT2 [25], and calculating BUSCO scores. Structural continuity was quantified by measuring gap percentages in 1 kb flanking regions of annotated genes.

### Transcriptome sequencing and annotation

Firstly, total RNA was extracted from brain, liver, spleen, and muscle tissues (TRIzol, Invitrogen), and high-quality samples (RIN ≥ 8.50) were sequenced on an Illumina HiSeq 4000 system (150 bp paired-end). Then, transcripts were assembled *de novo* using Trinity (v2.8.5) [26] to support annotation. The functional annotation pipeline integrated evidence from transcriptomics, homology, and *ab initio* predictions. Specifically, the Maker pipeline (v3.01.03) [27] combined evidence from the Trinity-assembled transcripts and homologous proteins from *Anas platyrhynchos* (SKLA 2.0), *Gallus gallus* (GRCg7b), *Mus musculus* (GRCm39), and *Homo sapiens* (GRCh38.p13). Concurrently, *ab initio* gene predictions were generated using GETA [28] and Funannotate (v1.8.9) [29]. A final non-redundant gene set was created by merging these sources, removing duplicates, and evaluating completeness using BUSCO (v5.2.2) [30] (aves_odb10). For haploid genomes, annotations were projected from the diploid reference using Liftoff (v1.6.3) [31].

### Phylogenomic analysis

For phylogenetic analysis, protein sequences from 13 species (including chicken, turkey, guinea pig, Pekin duck, Mallard duck, Muscovy duck, swan, human, mouse, zebra finch, lizard, platypus, dichromatic earthworm, blind earthworm, and tropical clawed frog) were clustered using OrthoMCL (v2.0.9) [32] and PANTHER (v17.0) [33]. Single-copy orthologs were aligned using Prank (v.170427) [34] (running 1,000 iterations under the "DNA" model for CDS sequences and the "AA" model for amino acid sequences), filtered by Gblocks (v0.91b) [35], and concatenated to construct a maximum likelihood tree using IQ-TREE (v1.6.5, JTT+G model) [36]. Optimal substitution models were selected based on the Bayesian Information Criterion (BIC). Subsequently, divergence times were estimated via TimeTree [37], and gene family expansion or contraction was analyzed using CAFÉ (v4.2.1) [38], and species trees were visualized using FigTree (v1.4.4) [39].

### Haplotype phasing and validation

Two complementary phasing strategies were applied to generate haplotype-resolved assemblies. The first approach utilized TrioCanu, where a contig-level phased assembly was generated using Canu (v2.2) [40] with 100x parental NGS and 120x ONT reads, followed by scaffolding against the diploid reference using RagTag (v2.1.0) [41].

The second approach employed Variation Graph (VG) phasing. In this pipeline, parental reads were mapped to the reference using BWA (v0.7.17) [42] and redundant alignments were removed using Samtools (v1.15) [43]. The non-redundant output was used for variant calling via Samtools. These parental variants were then encoded into parent-specific genome graphs where alternate haplotypes represented discrete paths. F1 sequencing reads were aligned to these graphs using vg giraffe (v1.40.0) [44], which utilized positional embeddings to ensure reads are mapped to the specific haplotype path consistent with their origin, thereby discriminating between paternal and maternal haplotypes based on path coverage differentials. This mapping information was used to generate VCF files containing the genotypes of F1 individuals. F1 reads were mapped to the graph (VG giraffe) to determine haplotype-specific genotypes, which were applied to the reference using BCFtools (v1.15) [45] to generate phased sequences. Phasing accuracy was evaluated using k-mer counting (Meryl and Merqury (v1.3) [46]) and collinearity analysis using MUMmer (v4.0) [47].

### Epigenomic and transcriptomic data analysis

Multi-omics data were processed to characterize haplotype-specific regulation. For chromatin architecture, clean Hi-C reads were mapped to phased genomes using BWA (v0.7.17) [60], separated by haplotype using a custom script, and analyzed for TADs, loops, and A/B compartments using the Juicer pipeline (v1.5) [21]. Hi-C data quality was rigorously controlled using HiC-Pro (v3.1.0) [48] to assess valid paired-end reads and interaction ratios.

For chromatin accessibility, ATAC-seq reads were aligned to haploid genomes using Bowtie2 (v2.4.5) [49]. Following alignment, Samtools (v1.15) was used for sorting and indexing, while Sambamba (v0.8.2) was employed to mark and remove PCR duplicates to ensure library complexity [43, 50]. Subsequently, bam files were converted to bed format using Bedtools (v2.30.0) [51], followed by peak calling using MACS2 (v2.2.7.1) [52]. ATAC-seq reproducibility was ensured using the Irreproducible Discovery Rate (IDR) framework (v2.0.4) [53] and Fraction of Reads in Peaks (FRiP) scores.

To assess allele-specific regulation, Allele-Specific Expression (ASE) and differential methylation were quantified. This analysis utilized SNPsplit (v0.5.0) [54] based on the phased SNP dataset to distinguish parental alleles in RNA-seq and bisulfite sequencing data, with allelic imbalance statistically identified by Qllelic (v0.3.2) [55] and nominal *P*-values were corrected for multiple testing using the Benjamini-Hochberg (BH) procedure across all tested genomic windows genome-wide to strictly control the false discovery rate (FDR). To ensure statistical reliability, only heterozygous sites supported by a minimum total read depth of 20 reads (with at least 2 reads per allele to minimize sequencing errors) were retained for analysis. A gene was defined as an ASE gene (demonstrating significant allelic imbalance) only if it met the following stringent criteria: a total allelic count greater than 20, a False Discovery Rate (FDR)-adjusted *P*-value of less than 0.05, and an absolute fold-change bias of log_2_(Paternal/Maternal) greater than 0.58 (corresponding to a >1.5-fold difference). Based on the consistency of this imbalance across biological replicates, ASE genes were further classified into deterministic monoallelic (DeMA) or random monoallelic (RaMA) categories as described in the Results.

### Identification of non-Mendelian inheritance regions

To systematically identify genomic regions deviating from Mendelian inheritance, we developed a comprehensive computational workflow integrating lineage tracing and multi-scale statistical screening. To distinguish the parental origin of F1 alleles, high-confidence parent-specific Single Nucleotide Variants (SNVs) were identified by comparing paternal and maternal whole-genome sequencing data against the diploid reference using GATK (v4.2.6.1) to serve as lineage markers [56]. F1 sequencing reads were aligned to the reference using BWA-MEM (v0.7.17) [42], and reads covering these parent-specific SNVs were rigorously categorized into paternal-derived reads containing only paternal-specific alleles, maternal-derived reads containing only maternal-specific alleles, or ambiguous reads containing shared and conflicting alleles. To strictly control for reference bias and sequencing errors, these ambiguous reads were explicitly excluded from downstream analysis. This conservative filtering strategy ensures that only high-confidence allele-specific assignments are used, preventing false-positive categorization arising from sequence similarity or mapping artifacts.

To detect non-Mendelian events ranging from local deviations to large-scale chromosomal anomalies, we implemented a variable sliding window strategy, scanning the genome with multiple window sizes of 100 kb, 500 kb, 1 Mb, 5 Mb, and 10 Mb to ensure robust detection across scales. Within each window, the raw aggregated counts of paternal and maternal reads were calculated and normalized by the total library size to correct for sequencing depth bias. Subsequently, the observed allelic ratio was tested against the expected 1:1 Mendelian segregation ratio using a Chi-square goodness-of-fit test. A genomic region was defined as a candidate non-Mendelian region only if it met stringent criteria, requiring that the deviation was statistically significant (*P*-value < 0.05) across overlapping windows, the region contained sufficient informative markers to ensure statistical power, and the direction of allelic bias remained consistent throughout the identified interval.

## Results

### High-quality diploid and haploid assembly

The genome size and heterozygosity of a female hybrid (CYF1-Female-1) from a Crimo male Muscovy duck (French) and a Yongchun female Muscovy duck (China) were estimated using the k-mer method (S1 Fig). We generated 110-fold standard Nanopore and 15-fold ultra-long Nanopore reads, 120-fold Bionano optical map molecules, 250-fold high-throughput chromosome conformation capture (Hi-C) reads, and 100-fold 150 bp Illumina paired-end reads of the CYF1, and 100-fold 150 bp Illumina paired-end reads from genomic DNA of its parents (S1-S7 Tables). Using a hierarchical and hybrid approach [57], we generated a high-quality diploid genome (*SKLFABB.CaiMos.1.0*), which spanned 1.23 Gb and is composed of 154 scaffolds with a contig N50 of 40.41 Mb, covering all chromosomes including 39 autosomes and two sex chromosomes (ZW) (Fig 1A-B, S8 Table).

**Fig 1.**
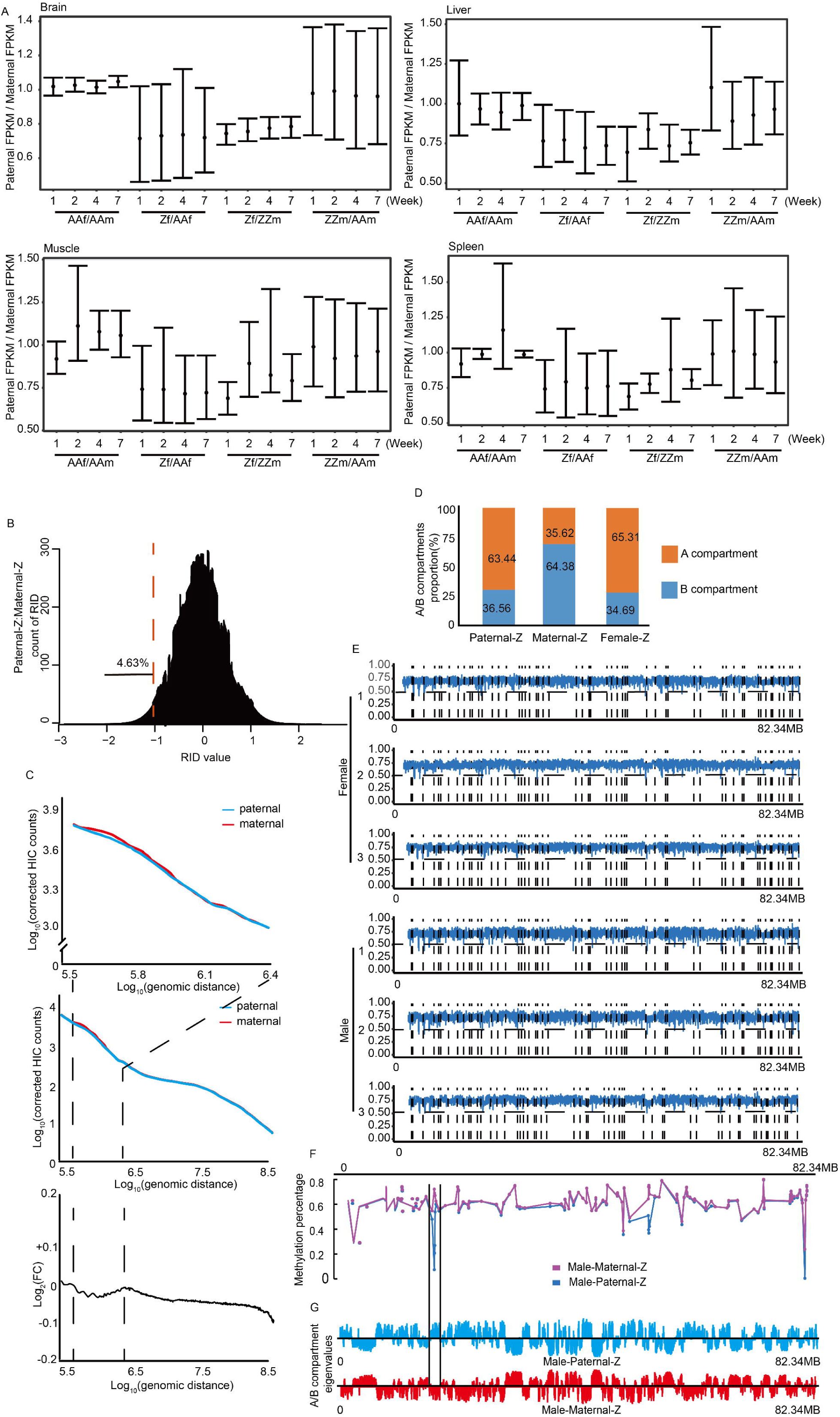
Assembly and evaluation of diploid and haploid Muscovy duck genomes. (A) Karyotype metrics of the diploid assembly showing chromosome size, GC content, and gene density. (B) Genomic feature distribution on representative macro- and micro-chromosomes (Chr1 and Chr29). (C) Workflow for generating paternal and maternal haploid assemblies. (D) Paternal (top) and maternal (bottom) synteny to the diploid reference (dark blue). Outer-to-inner tracks: phased regions (blue/brown), chromosomes (orange/red), and syntenic blocks (grey). (E) k-mer-based phasing completeness. Distributions of paternal-specific (red) and maternal-specific (blue) k-mers indicate clear haplotype separation. (F) Contiguity comparison with previous Muscovy duck, Pekin duck, and chicken genomes. (G) BUSCO completeness (%) for assemblies and proteomes. (H) Assembly integrity assessment via percentage of mapped RNA-seq reads. In all box plots, center lines represent medians, box limits indicate upper and lower quartiles, and whiskers extend to 1.5× interquartile range.

We then partitioned the 100-fold Nanopore long reads of the CYF1 into paternal and maternal subsets, and assembled each set into haploid contigs (Fig 1C). These haploid contigs were then independently scaffolded based on the diploid genome (*SKLFABB.CaiMos.1.0*), polished and filled gaps using haplotype-specific unique sequences. The final contig and scaffold N50 values after manual curation were 11.51 Mb and 81.82 Mb for the paternal assembly (*SKLFABB.CaiMos.1.0P*), and 10.52 Mb and 75.40 Mb for the maternal assembly (*SKLFABB.CaiMos.1.0M*) (S9 Table). Each haploid genome included 39 autosomes and one sex chromosome (Z or W), with 99.71% of paternal alleles and 99.02% of maternal alleles assigned to chromosomes, respectively (Fig 1D). Genome alignment and k-mer evaluation indicated that the haploid assemblies were highly syntenic to the diploid genome and clearly phased (Fig 1E).

Assessment of genome quality indicated that the diploid genome (*SKLFABB.CaiMos.1.0*) presented a significant improvement over the previous Muscovy genome (*KizCaiMos1.0* with a contig N50 of 11.8 Mb), higher than the chicken genome (*GRCg7b* with a contig N50 of 18.80 Mb) and comparable to that of our recently reported duck genome (*SKLA2.0* with a contig N50 of 32.90 Mb) in contiguity and completeness. The two haploid assemblies had 98.93% (*SKLFABB.CaiMos.1.0P*) and 98.65% (*SKLFABB.CaiMos.1.0M*) of the contigs longer than 1 Mb. Although they had slightly lower values than those of the diploid genome in contiguity, completeness and chromosome size, they were still comparable to those of the previous Muscovy duck genome (*KizCaiMos1.0*) and chicken (*GRCg7b*) (Fig 1F, S8-9 Tables). Further quality evaluation indicated that the diploid genome (*SKLFABB.CaiMos.1.0*) had a higher BUSCO score (Benchmarking Universal Single-Copy Orthologs, 96.7%), and higher mapping rates of population NGS reads (98.5-98.8%) and RNA-Seq reads (79.4-94.7%) than those of the previous genome (*KizCaiMos1.0*) (94.3% for BUSCO score, 98.2-98.5% for population NGS reads and 73.3-89.1% for RNA-Seq reads). Despite lacking one sex chromosome each, two haploid assemblies (*SKLFABB.CaiMos.1.0P* and *SKLFABB.CaiMos.1.0M*) had a similar level of BUSCO score (93.7, 93.9%), mapping rates of population NGS reads (97.4-98.1%) and RNA-Seq reads (79.13-93.66%) to the *KizCaiMos1.0* genome (Fig 1G-H). These results suggested that our new diploid and haploid Muscovy duck assemblies were high quality.

Subsequently, we predicted coding genes using Funannotate and the GETA pipeline with 175,405 assembled transcripts from RNA-Seq data and 108,127 full-length transcripts from PacBio sequencing (S10-11 Tables). This pipeline annotated 16,893 coding genes supported by 46,141 full-length transcripts for the diploid genome, 16,521 coding genes supported by 43,194 full-length transcripts for the paternal assembly, and 16,318 coding genes supported by 42,746 full-length transcripts for the maternal assembly. These Muscovy duck reference gene sets had 99.1%, 94.6%, 94.8% BUSCO score, 0.8%, 0.7%, 0.7% duplicate genes, 83.1-94.6% of RNA-seq mapping reads, and with 0.6%, 4.8%, 5.1% of genes missing. They are comparable to those in chicken (99.6%,0.2%, 86.6-95.7% and 0.3%) and duck (99.2%, 0.8%, 88.2-94.3% and 0.6%), but slightly superior to those of the previous Muscovy duck gene set (94.3%, 1.7%, 73.3-91.4% and 2.3%) (Fig 1G). These observations suggested that our Muscovy gene reference gene sets contain relatively complete gene sequences.

Moreover, we examined large-scale differences in gene complements between Muscovy duck and Pekin ducks using OrthoMCL [58]. This analysis revealed that the Muscovy duck genome significantly expanded 12 gene families and contracted 46 gene families when compared to the Pekin duck. Notable expansions occurred in olfactory receptor (*OR*) and protocadherin alpha-2 (*PCDHA*) gene families, while pronounced contractions characterized the very low-density lipoprotein receptor (*VLDLR*) and outer dense fiber protein (*ODF*) families, suggesting an adaptive specialization in sensory perception and lipid metabolism pathways (S12-13 Tables, S2 Fig).

### New efficient method for assembling haplotype genomes

We were inspired by pangenome graphs and adopted a similar graph-based pipeline to develop a new strategy to assemble haplotype genomes using short-read sequences from genomic DNA of both hybrid and parent (Fig 2A). This pipeline began by aligning parental short read sequences to the diploid genome (*SKLFABB.CaiMos.1.0*) and produced variant call format (VCF) files. These variant files were subsequently loaded into the variation graph (VG) toolkit [44] to construct parental-specific genome graphs encoding alternate haplotypes as discrete paths. Hybrid F1 reads were aligned to these graphs using positional embeddings, enabling haplotype discrimination through path coverage differentials. Path replacement with parental haplotypes ultimately produced phased VG-derived assemblies for both paternal and maternal genomes (Fig 2B-C).

**Fig 2.**
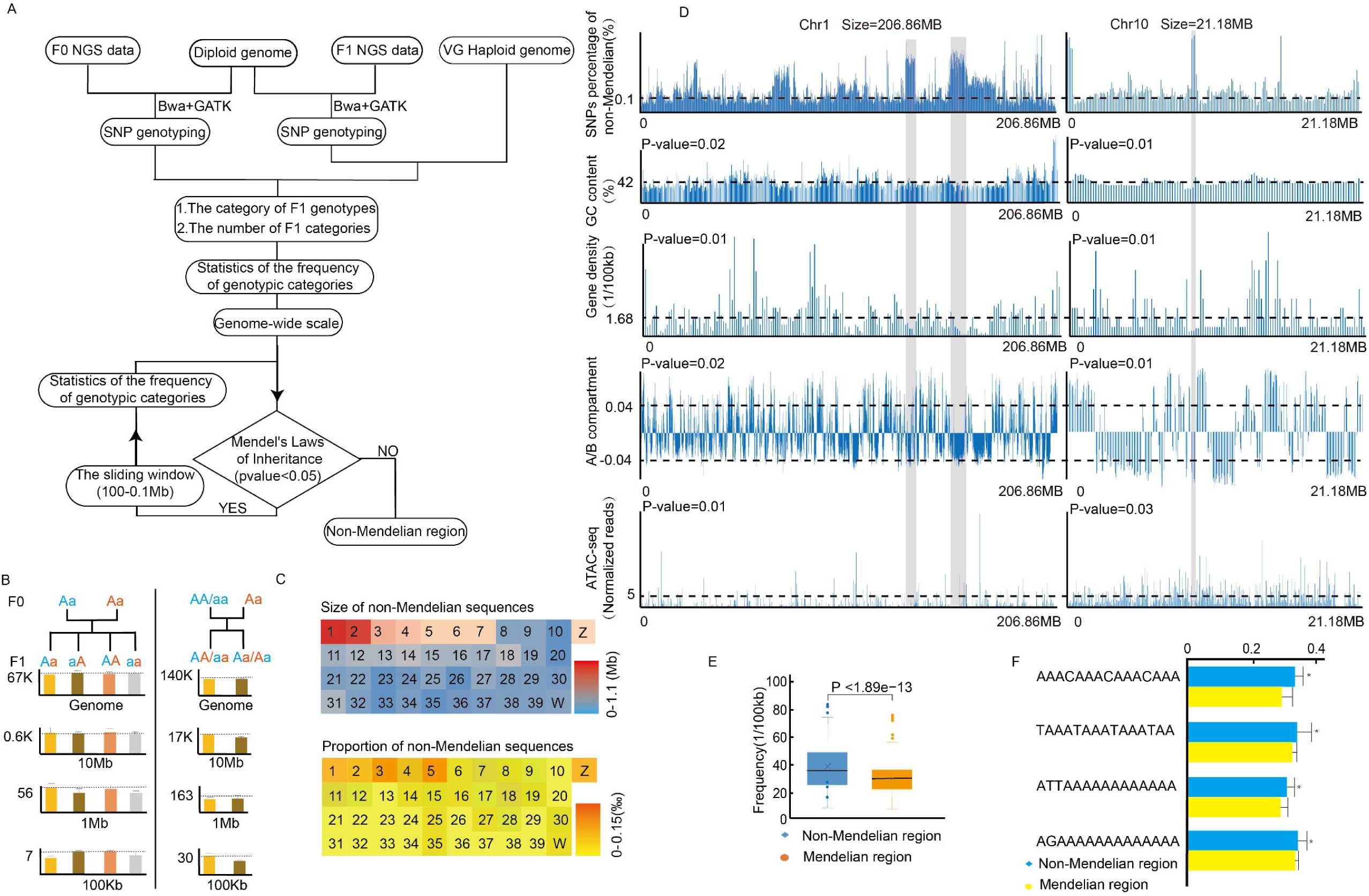
Construction of VG-haploid genomes and comparison with TrioCanu. (A) the Variation Graph (VG) phasing pipeline. (B) Paternal (top) and maternal (bottom) synteny to the diploid reference (dark blue) displaying outer-to-inner tracks of phased regions (blue/brown), chromosomes (orange/red), and syntenic blocks (grey). (C) k-mer-based phasing completeness evaluation. (D) Phased block size comparison with TrioCanu. (E) Genomic coverage comparison with TrioCanu. (F) Gene coverage comparison with TrioCanu. (G) BUSCO completeness (%) comparison with TrioCanu. (H) Assembly quality comparison with TrioCanu via percentage of mapped RNA-seq reads. In all box plots, center lines represent medians, box limits indicate upper and lower quartiles, and whiskers extend to 1.5× interquartile range. Abbreviations: Pat, paternal; Mat, maternal.

Using this new approach, we constructed 12 VG-haploid genomes for three full-sibling pairs of Muscovy ducks (three male and three female individuals) and predicted 16,800-17,100 protein coding genes (S3 Fig). The final contig and scaffold N50 values were 10-12 Mb and 75-82 Mb for the paternal VG-haploid genomes, and 10-12 Mb and 82.2-82.3 Mb for the maternal VG-haploid genomes, respectively (S14 Table). These were comparable to the corresponding values for paternal *SKLFABB.CaiMos.1.0P* (contig N50 of 11.51 Mb; scaffold N50 of 81.82 Mb) and maternal *SKLFABB.CaiMos.1.0M* (contig N50 of 10.52 Mb; scaffold N50 of 75.40 Mb) TrioCanu-haploid genomes. k-mer assessment indicated that these VG-haploid genomes were clearly phased similarly to TrioCanu-haploid genomes (S15 Table). Notably, although the VG-haploid genomes contained a lower total count of haplotype-specific k-mers (hapmers), they exhibited a 1.3-fold increase in hapmer block size, resulting in slightly larger phased blocks for both paternal and maternal genomes (Fig 2D). Moreover, the paternal VG-haploid genome had a genome size of 1.02 Gb with 15,458 genes being phased, and the maternal VG-haploid genome had 973 Mb with 15,360 genes being phased. These were slightly smaller than the corresponding values of the paternal (1.05 Gb and 15,773, *SKLFABB.CaiMos.1.0P*) and the maternal (991 Mb and 15,618, *SKLFABB.CaiMos.1.0M*) TrioCanu-haploid genomes (Fig 2E-F S16 Table).

In addition, the BUSCO assessment revealed high completeness for both haploid genomes and their reference gene set (S4 Fig). At the genome level, the VG-paternal haploid genome demonstrated 94.5% composite completeness (C), with 94.0% single-copy completeness (S), 0.5% duplicated (D), 1.1% fragmented (F), and 4.4% missing (M) orthologs. The VG-maternal haploid genome showed a composite completeness (94.3%), with a S of 93.8%, D of 0.5%, F of 1.0%, M of 4.7%. This was comparable to those of the TrioCanu-maternal and paternal genomes, where the former (*SKLFABB.CaiMos.1.0P*) had a 94.3% composite completeness (S: 93.9%; D: 0.4%; F: 1.1%; M: 4.5%), and the latter (*SKLFABB.CaiMos.1.0M*) displayed a 94.2% composite completeness (S: 93.7%; D: 0.5%; F: 0.9%; M: 4.9%). At the proteome-level, the VG-paternal coding genes had a 94.8% composite completeness (S: 94.1%; D: 0.7%; F: 0.4%; M: 4.8%), close to its genomic counterpart. The VG-maternal coding genes showed comparable metrics with 94.6% composite completeness (S: 93.9%; D: 0.7%; F: 0.3%; M: 5.1%). This was similar to that of the TrioCanu-paternal and maternal proteomes, where the former (*SKLFABB.CaiMos.1.0P*) had a 94.7% composite completeness (S: 94%; D: 0.7%; F: 0.4%; M: 4.9%), and the latter (*SKLFABB.CaiMos.1.0M*) had a 94.5% composite completeness (S: 93.8%; D: 0.7%; F: 0.4%; M: 5.1%) (Fig 2G, S17 Table). Additionally, the VG-haploid genomes had similar RNA-seq mapping reads (85.62-92.91% for paternal and 81.13-93.66% for maternal VG-haploid genomes) to those of TrioCanu-haploid genomes (83.71-93.66% for paternal and 82.17-92.14% for maternal VG-haploid genomes) (Fig 2H), indicating this new approach could assemble haploid genomes with high quality. Genome alignment and k-mer evaluation indicated that the haploid assemblies were highly syntenic to the diploid genome and clearly phased (S5 Fig).

### Paternal and maternal alleles expression patterns

Using RNA-seq data of three full-sibling pairs of Muscovy ducks, we identified 1,840,361 heterozygous single-nucleotide polymorphisms (SNPs), which covered more than 75% of genes (13,091 genes for paternal haploid genomes and 12,863 genes for maternal haploid genomes). We phased reads based on SNP information and characterized allelic expression patterns by aligning RNA-Seq reads of four Muscovy duck tissues (brain, liver, spleen, and muscle) to three full-sibling pairs haplotype genomes. In general, maternal alleles exhibited a similar number of expressed genes (9,024-10,964) but showed relatively higher expression levels (MAT:PAT = 1.05, 1.03, 1.05, 1.10) when compared to paternal alleles (brain: 11,055; liver: 8,825; spleen: 10,548; muscle: 10,603; Fig 3A).

**Fig 3.**
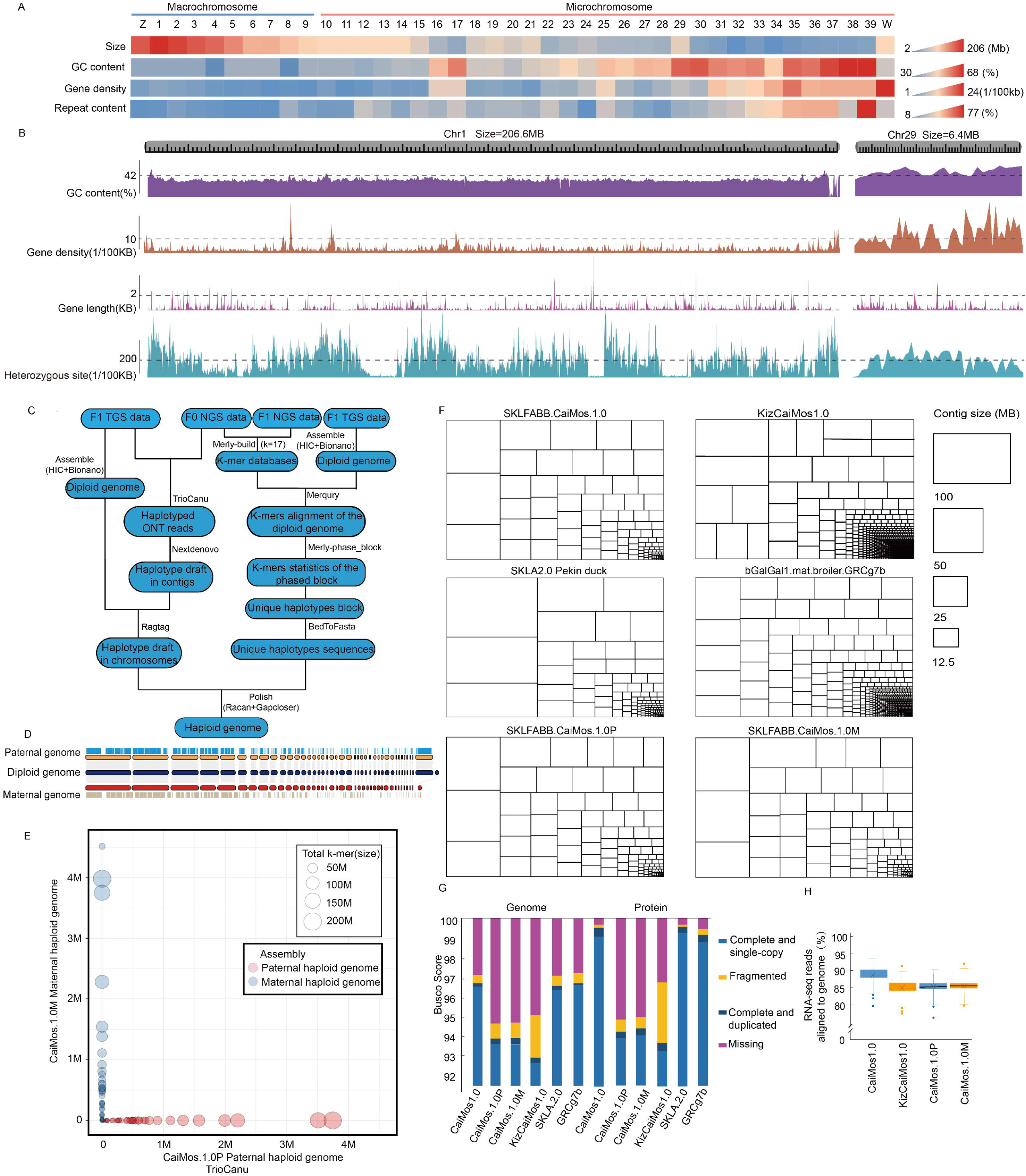
Paternal and maternal gene expression patterns in Muscovy duck tissues. (A) Density plot of log_10_(Paternal/Maternal) expression ratios across brain, liver, spleen, and muscle, where the dashed line indicates balanced expression. (B) Plots of paternal fraction highlighting genes with significant paternal or maternal bias (red). The y-axis represents the paternal fraction (paternal reads/total reads), centered at 0.5. (C) Venn diagram showing the overlap of shared and tissue-specific paternally and maternally biased genes.

Among 8,825-11,055 expressed genes in brain, liver, spleen and muscle tissues, 15.49-17.77% (1,712-1,744), 27.24-30.13% (2,404-3,441) and 52.38-52.99% (4,677-5,901) showed biallelic, deterministic monoallelic (DeMA – the allele selection is predetermined, 10.90-12.45% for paternal-mono and 16.34-18.68% for maternal-mono), and biased/random autosomal monoallelic (RaMA – the allele selection is not predetermined and each allele has an equal chance to become the active or inactive allele) states, respectively (Fig 3B). Detailed analysis revealed that while the majority of DeMA genes showed tissue-specific patterns, a small subset of 5.19% (125 genes) was conserved across tissues. These conserved DeMA genes were enriched in essential pathways, including Oxidative phosphorylation, Protein biosynthesis, and RNA splicing machinery (*FDR* < 0.05) (S6 Fig). Specifically, five genes (*ALAS1, EIF3L*, *PRCC, SEPTIN6* and *SYAP1*) were mat-mono (deterministic maternal) genes and three genes (*MOSPD1, CSRNP1* and *SRP54*) were pat-mono (deterministic paternal) in the above four tissues (Fig 3C). When compared to DeMAs, 4,677 RaMAs displayed higher tissue specificity, where muscle showed the highest proportion (3,189 genes), and only 48 (1.03%) genes involved in antigen presentation, olfactory receptor activity, and protocadherin clustering, exhibited cross-tissue conservation.

### Paternal and maternal chromatin conformation landscapes

Using the above 12 haploid genomes of three full-sibling pairs of Muscovy ducks, we characterized parent-of-origin pattern on chromatin structure using phased Hi-C data (Fig 4A, S7-8 Figs, S18 Table). At the genome-wide level, paternal haplotypes displayed a more compact chromatin architecture, with 5.13% interchromosomal interaction bins showing significantly higher RID (Relative Interaction Difference) values (> 1) when compared to maternal haploid genomes (Fig 4B). Paternal haploid genomes had a slight decrease in short-distance chromatin contact and a global increase in long-distance chromatin contact (Fig 4C-D). Contact decay analysis showed a 7.8% lower frequency of short-range (≤1 Mb) contacts and 12.3% higher frequency of long-range (>10 Mb) contacts in paternal haplotypes (*P* = 0.004, n = 3), accompanied by a shallower decay exponent (α = -1.10 vs -1.16).

**Fig 4.**
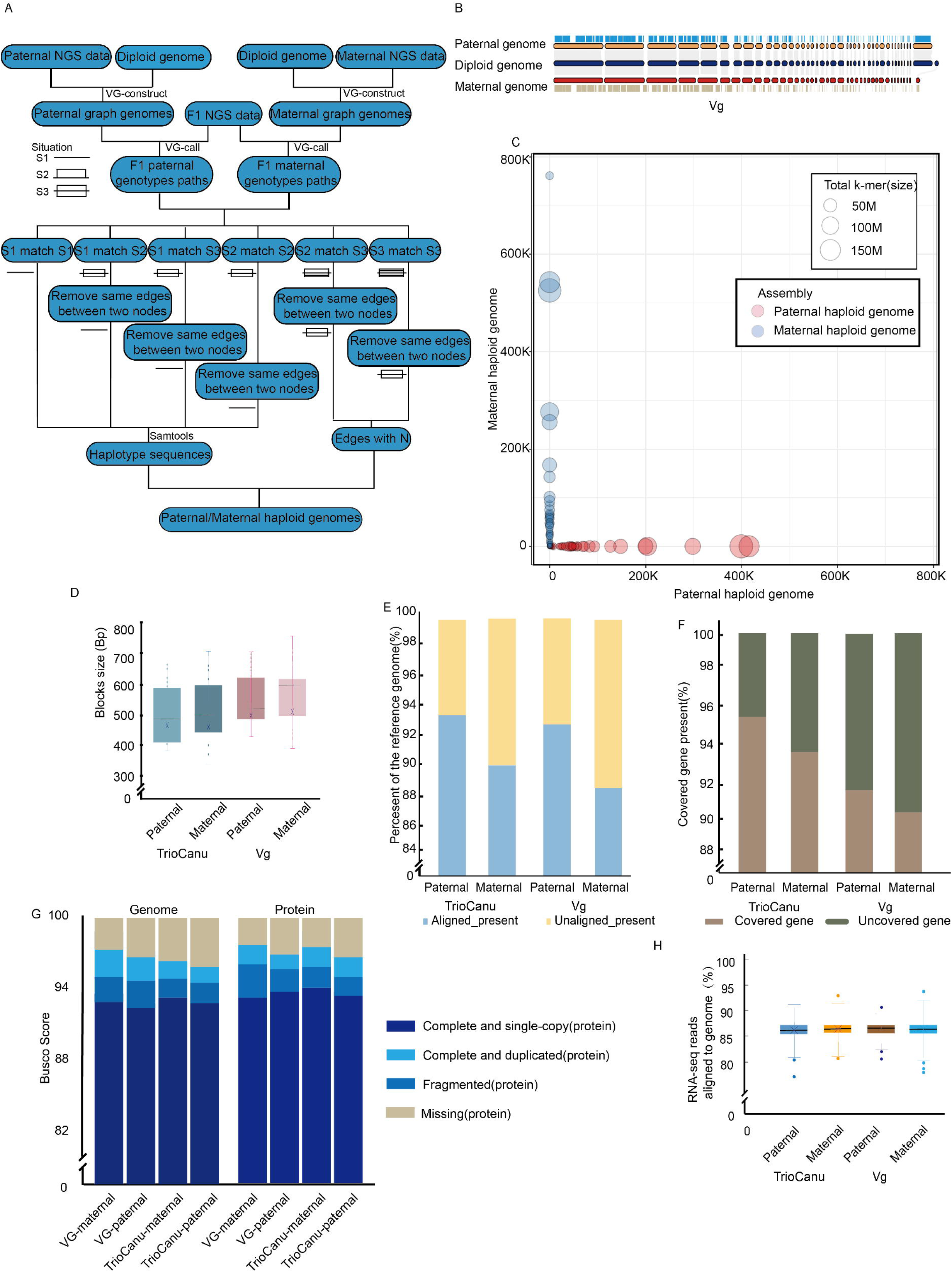
Paternal and maternal chromatin architecture differences in Muscovy duck. (A) Heatmap of differential inter-chromosomal interaction frequencies calculated as paternal minus maternal. (B) Distribution of Relative Interaction Difference (RID) values where red dashed lines indicate the significance cutoff (RID > 1 or < -1, representing a two-fold difference). (C) Proportion of differential interactions biased toward paternal or maternal haplotypes. (D) Intra-chromosomal interaction decay showing contact frequencies and log2 fold changes versus genomic distance. (E) Genome-wide A/B compartment proportions and inter-haplotype stability. (F) Comparison of compartment number (top) and length (bottom) between haplotypes. (G–K) Paternal and maternal comparison of compartment strength (G), TAD number (H), TAD length (I), intra-TAD interaction strength (J), and TAD boundary strength (K). In all box plots, center lines represent medians, box limits indicate upper and lower quartiles, and whiskers extend to 1.5× interquartile range.

At the functional compartment scale, we identified 582 compartments (median size = 2.06 Mb, gene coverage = 99.13%) for paternal haploid genomes, and 570 compartments (median size = 2.11 Mb, gene coverage = 99.16%) for maternal haploid genomes (S9-10 Figs). When compared to maternal haploid genomes containing 48.98% A (active) compartments and 51.02% B (inactive) compartments, paternal haploid genomes had a slightly lower proportion (48.89%) of active (A) compartments but a higher proportion (51.11%) of inactive (B) compartments (Fig 4E, S11 Fig). Detailed analyses indicated that most (99.91%) regions of parental haploid genomes were in the same (A or B) functional compartments, and a small proportion (1.07 Mb, 0.09%) of parental haploid genomes were segregated into opposite functional compartments (one was in the A compartment and the other was in the B compartment) (Fig 4E). These parental A/B switched regions were enriched in genes related to the regulation of phosphoprotein phosphatase activity (FDR = 3.2×10[[), the Hedgehog signaling pathway (FDR = 1.8×10[³) and the TGF-β signaling pathway (FDR = 4.1×10[³) (S12 Fig). Among them, 0.61 Mb of paternal haploid genomes in A compartment contained the *RPRD2* and *P4HB* genes, while 0.46 Mb of maternal haploid genomes in A compartment contained the *ZNF598* and *USP22* genes.

We further examined interaction within compartments and detected significant differences in compartment strength between paternal and maternal haploid genomes. We found that paternal and maternal haploid genomes generally showed similar levels of interaction within compartments and compartment strength, but had 7.24 Mb of A/B variable regions presenting differences in interactions within compartments (Fig 4F-G, *P* = 0.006). These A/B variable regions contained genes enriched in the RNA degradation (FDR = 6.7×10[[), WNT signaling pathway (FDR = 2.9×10[³), and metabolic pathways (FDR = 7.1×10[³) (S13 Fig). Among them, 3.72 Mb of paternal haploid genome showed higher interaction strength within compartments containing the *WNT5A* and *WNT7B* genes, while 3.48 Mb of maternal haploid genomes had higher interaction strength within compartments containing the *TBL3* gene.

Subsequently, we compared TAD (topologically associated domain) features of parental haploid genomes in the hybrids. We observed that paternal and maternal haploid genomes had the same number (1612) of TADs with a median size of 0.74 Mb. Both parental genomes showed no difference in interaction strength of TADs (median in paternal = 0.72, median in maternal = 0.72, *P* = 0.82), position of boundaries and boundary strength (median in paternal = 3.22, median in maternal = 3.21, *P* = 0.76, Fig 4H-K and S14 Fig).

### Paternal and maternal epigenetic landscapes

We compared paternal and maternal DNA methylation (DNAm) levels using phased whole-genome bisulfite sequencing data of muscle and brain from three full-sibling pairs of Muscovy ducks. In general, paternal haploid genomes had similar methylation profiles (98.59%) of DNAm regions to those of maternal haploid genomes (98.55%) (Fig 5A). We assayed DNAm levels at the gene level, and separately for CpG islands which were enriched in promoter regions. As expected, gene methylation levels were high (mean ± SD: 65.3% ± 6.7%), substantially lower in regulatory CpG islands (0.65% ± 1.10%) and near-zero within ATAC-seq peaks (0.63% ± 0.30%) in both parental haploid genomes. At the gene level, paternal and maternal haploid genomes showed similar DNAm levels in brain and muscle tissues. In CpG islands, paternal and maternal haploid genomes presented significant differences in DNAm landscapes at 158 genes.

**Fig 5.**
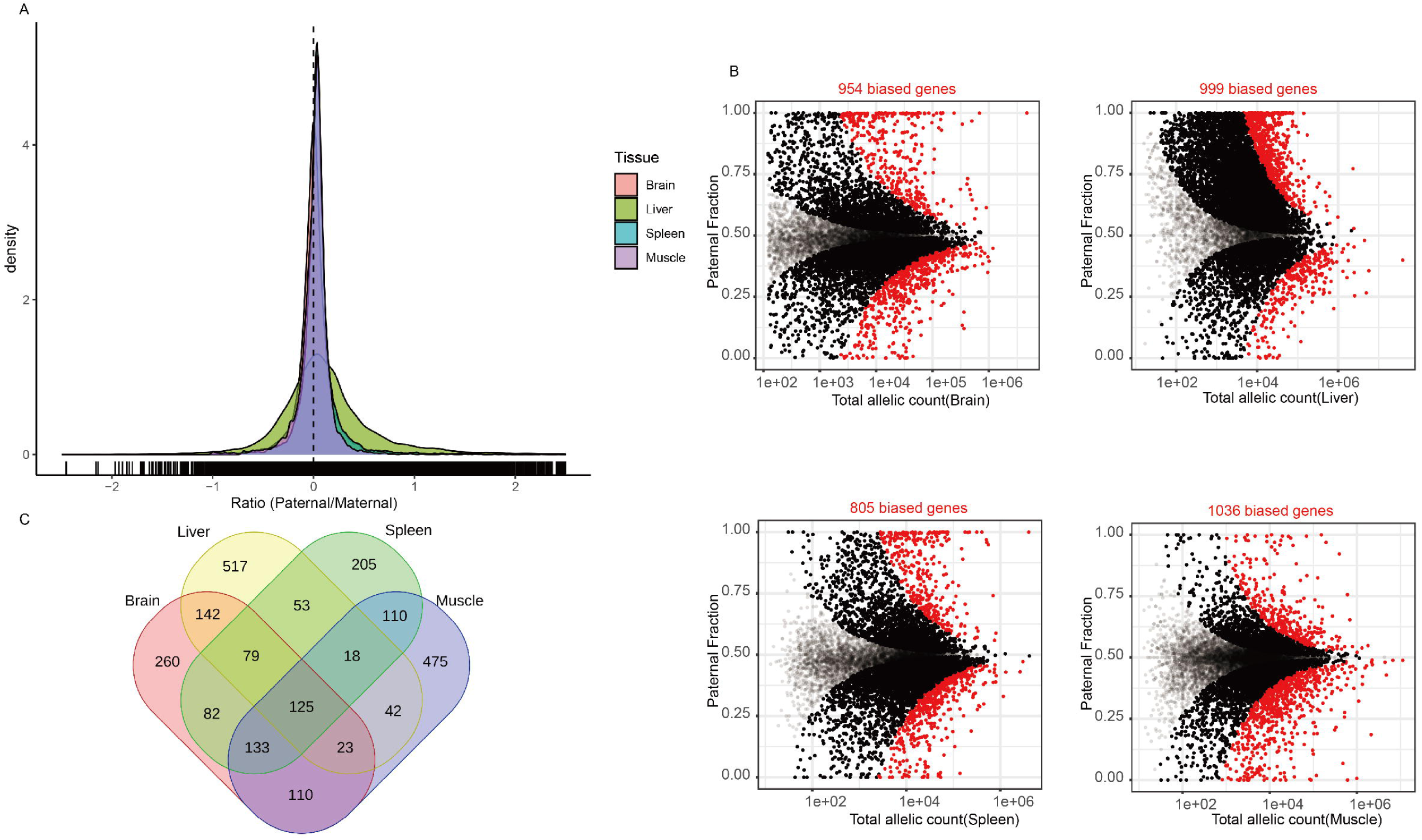
Paternal and maternal methylation and chromatin accessibility landscapes. (A) Genome-wide distribution of paternal-to-maternal methylation ratios across autosomes for female and male individuals. (B) KEGG pathway enrichment analysis of differentially methylated genes. (C) Comparison of ATAC-seq peak heights and signal distribution along Chromosome 1 between paternal and maternal haplotypes. (D) Heatmap showing paternal-maternal chromatin accessibility differences at transcription start sites. (E) Venn diagram of total ATAC-seq peaks identified on paternal and maternal haplotypes. (F) Venn diagram of genes associated with ATAC-seq peaks on paternal and maternal haplotypes. In all box plots, center lines represent medians, box limits indicate upper and lower quartiles, and whiskers extend to 1.5× interquartile range.

We further categorized methylated regions into 74 paternal allele-specific DNAm genes/CpG islands (Ps-DNAms), 50 maternal allele-specific DNAm genes/CpG islands (Ms-DNAms) and 34 biallelic DNAm genes/CpG islands (Bi-DNAms). Ms-DNAms were enriched in TSSs (Transcription Start Sites) of genes related to protein transmembrane transport (*SRP54*), RNA splicing regulation (*SRSF5*), and mitochondrial metabolic pathways (*DLST*), and included pat-mono genes such as *MOSPD1* and *CSRNP1*. In contrast, Ps-DNAms were enriched in TSSs of genes related to cellular stress response (*EIF3L*), heme biosynthesis (*ALAS1*), and cytoskeletal organization (*SEPTIN6*), and included mat-mono genes (*SYAP1*) involved in maternal allele-specific regulation (Fig 5B). For Bi-DNAms, we observed that hypomethylated regions (≤5%) over TSSs of paternally expressed genes including *SRSF5* and *DLST* were hypermethylated (≥20%) in maternal haploid genomes, suggesting parent-of-origin-dependent epigenetic silencing. In contrast, regions over TSSs of maternally expressed genes including *SEPTIN6* and *EIF3L* were hypermethylated (≥20%) in paternal haploid genomes but hypomethylated (≤5%) in maternal haploid genomes, indicating allele-specific methylation dynamics.

Subsequently, we profiled paternal and maternal haploid genomes chromatin accessibility landscapes using phased ATAC-seq data of three full-sibling pairs of Muscovy ducks (S19 Table). This detected a global atlas of 59,266 chromatin peaks covering 12,788 TSS sites for paternal genomes and 59,563 chromatin peaks covering 13,007 TSS sites for maternal genomes (Fig 5C-D). In general, paternal and maternal haploid genomes exhibited significant differences in peak height at accessible chromatin sites near transcription start sites (TSSs, *P* = 0.034), suggesting allele-specific variation in chromatin organization. Further comparison identified 1125 paternal allele-specific ATAC-seq peaks (Ps-ATACPs), 1422 maternal allele-specific ATAC-seq peaks (Ms-ATACPs) and 58,141 biallelic ATAC-seq peaks (Bi-ATACPs, Fig 5E-F). Ps-ATACPs were enriched in TSSs of *EIF1AX, HARS1, LHX3, LOC113842949, LOC119713031, LRRC14*, while Ms-ATACPs were enriched in TSSs of *E2F4, GPR27, LOC113842963, PENK*. For Bi-ATACPs, we observed that 8,305 peaks were significantly elevated in paternal haploid genomes covering 1,766 TSSs of 1,964 genes, whereas 29,113 peaks showed maternal haplotype-specific elevation spanning 6,302 TSSs of 6,879 genes. We integrated the ATAC-seq data with the above RNA-seq and whole-genome bisulfite sequencing data. We found that TSSs of 10 genes including *E2F4, GPR27, LOC113842963, PENK, EIF1AX, HARS1, LHX3, LOC113842949, LOC119713031* and *LRRC14* showed significant differences in chromatin accessibility (ΔATAC > 2-fold), methylation levels (|Δβ| > 0.3), and expression (|log_2_FC| > 1) between the paternal and maternal alleles (S15 Fig). These observations might provide insights into parental allele-specific epigenetic regulation and gene expression mechanisms in Muscovy duck, particularly through the coordinated interplay of DNA methylation, chromatin accessibility, and transcriptional activity at loci such as *E2F4, GPR27, LHX3,* and *PENK*.

### Paternal and maternal heterogeneity of Z activation

Using 12 haploid genomes of three full-sibling pairs of Muscovy ducks, we characterized the Z chromosome activation at multiple scales. Firstly, we evaluated gene expression levels on a chromosome-wide scale. RNA-seq data of muscle and brain from Muscovy ducks at week 1, 2, 4 and 7 demonstrated partial dosage compensation (0.5 < Z_f_ [female Z chromosome] : AA_f_ [female autosomes] < 1, AA_f_ : AA_m_ [male autosomes] ≌ 1.05, Z_f_ : AA_f_ ≌ 0.74, ZZ_m_ [male Z chromosome] : AA_m_ ≌ 0.92) and dosage balance (0.5 < Z_f_ : ZZ_m_ < 1, Z_f_ : ZZ_m_ ≌ 0.89, ZZ_m_ : AA_m_ ≌ 0.92) on the Z chromosome through the regulation of Z-linked gene expression in females and males (Fig 6A, S16 Fig). This is consistent with gene expression patterns of the Z chromosome in chicken and crow [59–61]. We then characterized gene expression pattern of the paternal and maternal Z chromosomes (Zp and Zm) using phased RNA-Seq data. In general, the Zp of females had similar numbers of expressed genes (584-620), but exhibited higher expression levels (525-550) than both the Zp and Zm of males in all four tissues (brain, liver, spleen and muscle) (Fig 6A).

**Fig 6.**
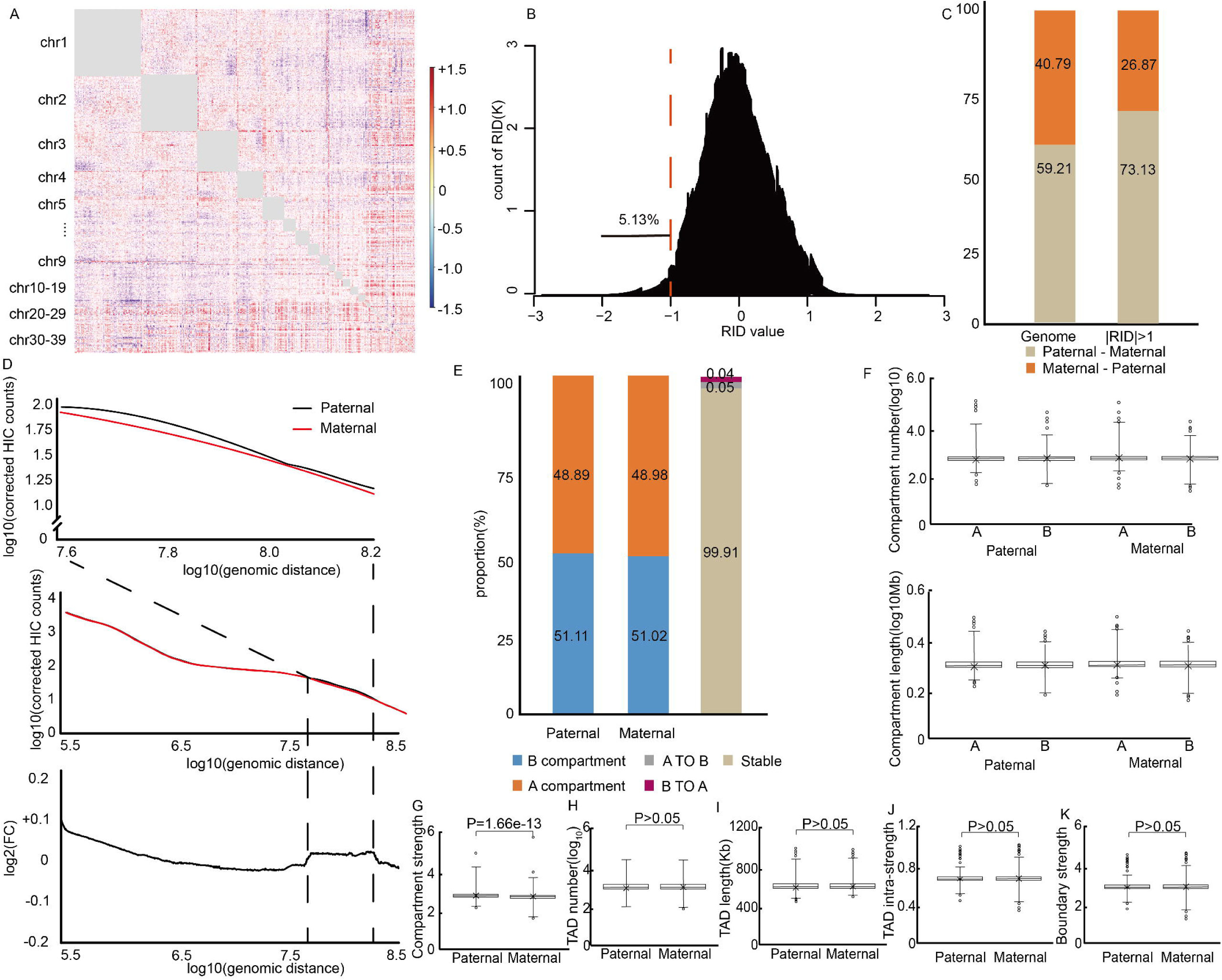
Paternal and maternal expression, chromatin architecture, and methylation on the Z chromosome. (A) Paternal-to-maternal expression ratios across developmental stages in brain, liver, muscle, and spleen. (B) Distribution of Relative Interaction Difference (RID) values where the red dashed line indicates the significance cutoff. (C) Intra-chromosomal interaction decay comparing paternal and maternal Z chromosomes. (D) Proportions of A and B compartments on paternal, maternal, and female Z haplotypes. (E) Methylation percentage across the Z chromosome. (F) Detailed view of paternal and maternal methylation ratios. (G) A/B compartment eigenvalues for paternal and maternal Z chromosomes.

Secondly, we compared chromatin structure of the Zp and Zm using phased Hi-C data of the above three full-sibling pairs of Muscovy ducks. We calculated relative interaction difference (RID) between the Zp of females, the Zp and Zm of males. We found interchromosomal interactions were significantly decreased in the female Zp (*P =* 4.3×10^-9^), and slightly reduced in the male Zp (*P =* 0.017) when compared to the male Zm. Among 809 interchromosomal interaction bins detected on the Zp and Zm, the numbers of slightly lower RID values (< 0) and significantly lower RID values (< -1) were 389 (48.1%) and 38 (4.71%) for the female Zp, and 398 (49.2%) and 37 (4.63%) for the male Zp when compared to the male Zm (Fig 6B-C, S17 Fig). At the functional compartment scale, the female Zp had the highest proportion (65.31%) of the A compartment and the strongest compartment strength, the male Zp had a higher proportion (63.44%) of the A compartment and the stronger compartment strength, and the male Zm had a low proportion (35.62%) of the A compartment and the weak compartment strength (Fig 6D, S18 Fig, S20 Table). This is consistent with the observation that the female Zp contained the largest number and highest levels of expressed genes, while the male Zm had the smallest number and lowest levels of expressed genes. This was supported by gene expression on the Z chromosome showing significantly higher connection strength to interchromosomal interaction than that on autosomes. Surprisingly, a large proportion (96.63%) of the male Zp and Zm regions showed an opposite compartment status of the A/B compartment, where one parental allele was in the A compartment and the other one was in the B compartment. These regions contained 36.5% (148/406) genes on Z chromosomes, which showed parental allele-specific expression and were enriched in cytokine-related processes (FDR *=* 3.6×10^-10^), including cytokine-mediated signaling pathways (FDR *=* 3.8×10^-8^) and cellular responses to cytokine stimulus (FDR *=* 1.2×10^-5^) (S19 Fig). In contrast, only a low proportion (0.9%) of parental allele-specific expressed genes demonstrated opposite distribution of A/B compartments on autosomes (Fig 4E). At the topologically associating domain (TAD) scale, we found that two parental Z chromosomes were conserved, which included the same number of TADs (123), TAD size (613.18 ± 28.45 kb) and covered genes (760-762), and did not change boundary strength (*P* = 0.64) (S20 Fig, S21 Table).

Thirdly, we profiled DNA methylation and chromatin accessibility map of the Zp and Zm using phased data of the above three full-sibling pairs of Muscovy ducks. Concordant with gene expression and chromatin structure pattern, the female Zp displayed the lowest methylation level (*P* = 1.7×10^-5^) and highest chromatin accessibility; the male Zp had an intermediate methylation level and chromatin accessibility; and the male Zm showed the highest methylation level and lowest chromatin accessibility (ΔATAC = 0.12 ± 0.15, *P* = 0.02) (Fig 6E-G). However, we only identified 7 ASE genes with allelic differential methylation (*CENPH, LINGO2, LOC101792280, LOC101792981, LOC121108514, MLANA, MTREX*) showing significant parental bias (*P* < 0.01). Ms-DNAms covered *CENPH, LINGO2, LOC101792280,* and Ps-ATACPs covered *LOC101792981, LOC121108514, MLANA, MTREX*. Based on the observations, it appears that methylation is not the primary cause of allelic imbalance in the Z chromosome genes at these developmental stages.

At last, we compared chromatin accessibility profiling of paternal and maternal haploid genomes. This identified 217 paternal allele-specific ATAC-seq peaks (Ps-ATACPs), 156 maternal allele-specific ATAC-seq peaks (Ms-ATACPs), and 3,349 biallelic ATAC-seq peaks (Bi-ATACPs). Ps-ATACPs were predominantly enriched near transcription start sites (TSSs) of genes associated with cytokine-cytokine receptor interactions and the NF-κB signaling pathway, whereas Ms-ATACPs primarily localized to TSS regions of genes regulating the mTOR signaling pathway and cell cycle progression. Among the 3,349 Bi-ATACPs, allele-specific chromatin accessibility dynamics were observed: 573 peaks (spanning 673 genes and covering 635 TSSs) exhibited significantly elevated accessibility in paternal haploid genomes, while 162 peaks (spanning 139 genes and covering 108 TSSs) showed enhanced accessibility in maternal haploid genomes (S21 Fig, S22 Table). This analysis identified 40 genes with allelically biased TSS peak clustering, including TSSs of two genes (*LOC112531540* and *MFSD5*) displayed exclusive paternal peak clustering on the Z chromosome and TSSs of two genes (*LOC101750948* and *SCAMP1*) demonstrated maternal-specific peak clustering (S22 Fig).

### Identification of non-Mendelian regions using haploid genomes

We evaluated genetic inheritance in the whole genome based on genotypes of parents and offspring (6 F0 and 6 F1 individuals) using 12 haplotype genomes (Fig 7A-B). This evaluation was carried out as the following pipeline. First, F1 alleles were genotyped by aligning NGS reads to their respective haploid genomes. Next, the parental origin of these F1 alleles was traced by aligning parental NGS reads to the diploid reference genome (*SKLFABB.CaiMos.1.0*). Subsequently, we quantified the frequency of various genotypic categories, incorporating parent-of-origin information for the F1 individuals. The genome was then segmented using sliding windows ranging from 100 kb to 10,000 kb to categorize genotypes within each window. (S23 Fig), Finally, potential biases in Mendel’s laws were evaluated by assessing the frequency of genotypic categories across all windows.

**Fig 7.**
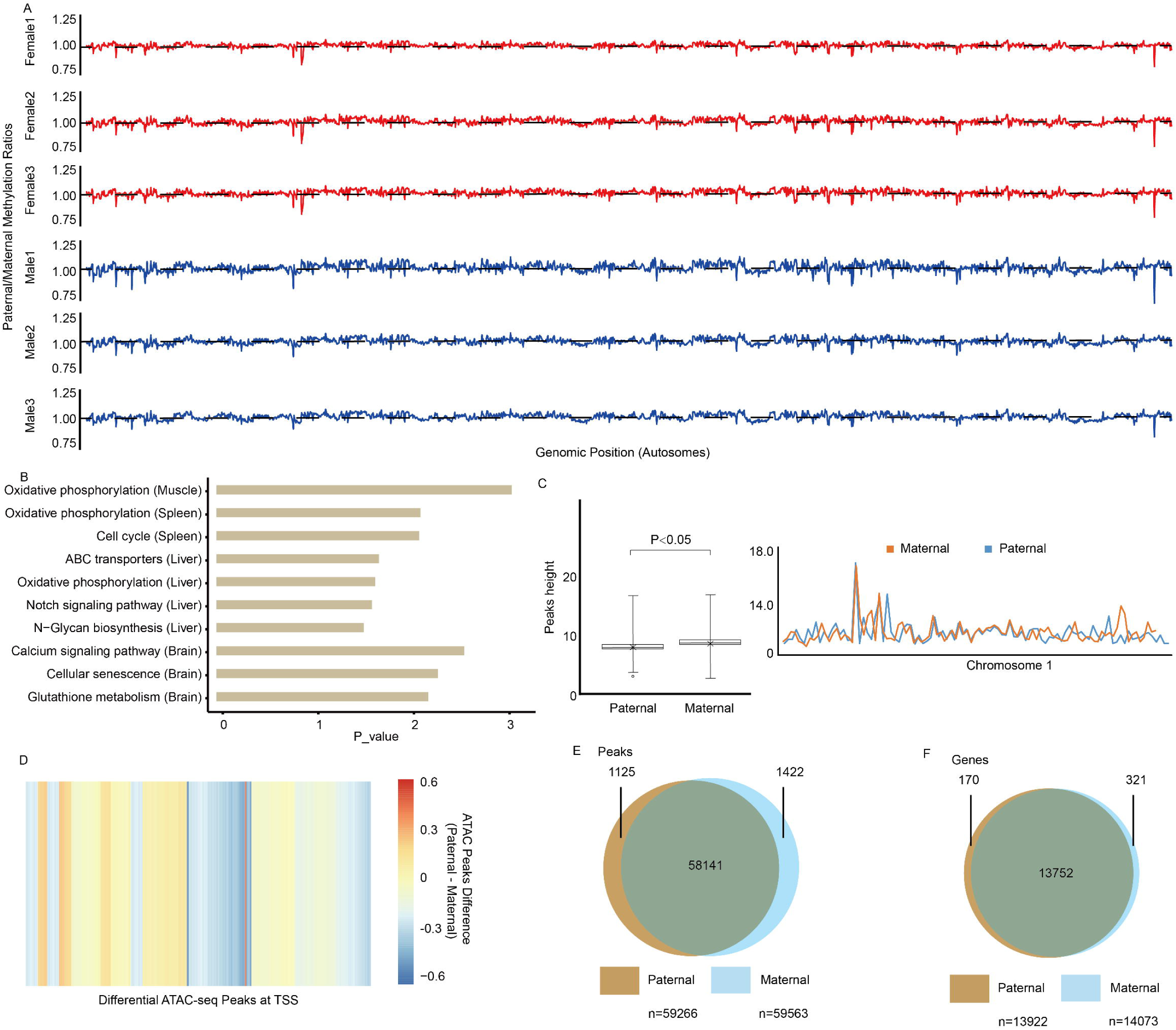
Detection and characterization of non-Mendelian inheritance regions. (A) Pipeline for identifying non-Mendelian regions from parental and F1 NGS data using multi-scale statistical screening. (B) Counts of genotypic categories detected using different sliding window sizes. (C) Heatmap displaying the distribution of non-Mendelian regions across chromosomes. (D) Genomic and epigenomic landscape comparison between non-Mendelian (grey shaded) and Mendelian regions on Chromosomes 1 and 10. (E) Comparison of repetitive sequence frequency in non-Mendelian and Mendelian regions. (F) Box plots define the median (center line), upper and lower quartiles (box limits), and 1.5× interquartile range (whiskers). Significance was determined by the Wilcoxon rank-sum test (*P* < 0.05).

This effort found that 549,387 genotyped loci followed a 1:1 segregation ratio when they were scanned under 10 Mb windows, each including about 700 genotyped loci. However, 1,428 of 549,387 (0.26%) genotyped loci deviated from the expected 1:1 segregation ratio when they were scanned under 1 Mb windows, each including about 130 genotyped loci (Fig 7B). These non-Mendelian regions covered 3.18 Mb and were enriched on chromosomes 1-5, where chromosomes 1-5 occupied 56.17% of the genome, but contained 86% of the non-Mendelian regions (Fig 7C).

We characterized these non-Mendelian regions at multiple scales. Firstly, we found that in sequence composition, the non-Mendelian regions had a similar SNP density (0.1027 vs 0.1025 SNPs/100 bp), but significantly lower gene density (1.19 vs 3.24 genes/Mb), GC content (38.41% vs 41.62%) and repetitive sequence frequency (30.06 vs 38.75 counts/100 kb) compared to those of the Mendelian regions (Fig 7D-E, S24 Fig). Subsequently, we scanned motifs and found that the non-Mendelian regions were significantly enriched for four types of DNA motifs (AAACAAACAAACAA, TAAATAAATAAATAA, ATTAAAAAAAAAAAA and AGAAAAAAAAAAAAA; Fig 7F, S23 Table). Syntenic analyses demonstrated that the above four types of DNA motifs were enriched in homologous regions of non-Mendelian regions from five divergent avian genomes (Chicken, Turkey, Mallard, Pekin duck and Kiwi; S25-26 Figs). Previous studies indicated that these DNA binding motifs were bound by the forkhead family of transcription factors including FOXC1, FOXP1 and FOXD3. Such binding between the DNA motifs and these forkhead box proteins promotes DNA bending, thus might affect chromatin structure. We then compared chromatin structures of the non-Mendelian and Mendelian regions. Surprisingly, the non-Mendelian regions had a more compact chromatin organization with a predominantly higher proportion (86.11% vs 50.17%) of regions in the B compartments and lower proportion (13.89% vs 49.83%) of regions in the A compartments when compared to the Mendelian regions (Fig 7D). This relatively compacted chromatin organization of non-Mendelian regions was consistent with their low chromatin accessibility, where the non-Mendelian regions had a lower average peak value (2.61 vs 5.38) of ATAC-seq when compared to the Mendelian regions (Fig 7D). These observations suggested that low GC content and low repeat sequence frequency, especially the enriched DNA motifs might affect chromatin organization and accessibility, thus influencing allele segregation during meiosis and contribute to the appearance of non-Mendelian regions.

## Discussion

Hybrid breeding is a revolutionary technology widely adopted in agriculture to increase productivity by generating heterosis. Hybrid breeding also generates hybrid dysgenesis showing phenotypic abnormalities. Thus, it is important to understand the biological mechanism for heterosis, design hybrid breeding to generate heterosis and avoid hybrid dysgenesis in agriculture. Haplotype-resolved genome assemblies provide important resources to understand how expression and regulation of parental alleles impact heterosis [62–65]. However, current haploid genome assemblies rely on long-read sequencing technology producing reads that are tens of kilobases long, such as PacBio SMRT sequencing and Oxford Nanopore sequencing [66–69]. Due to the throughput and cost of long-read sequencing technology, this technology is limited to estimate genetic, expression and regulatory patterns of parental alleles in hybrid breeding with large populations. Here we developed an efficient and cost-effective method to assemble haploid genomes using high throughput short-reads of hybrid and parents. Using this new method, we constructed 12 VG-haploid genomes, which were clearly phased and showed similar quality in genomic contiguity and completeness to that of haploid genomes assembled with phased Nanopore reads (Fig 2G-H). Although these VG-haploid genomes only distinguished ∼75% of paternal and maternal alleles due to limited heterozygous SNPs (1,840,361), we successfully characterized parental patterns in genetics, expression and regulation. These observations suggested the possible utilization of our method in understanding the biological mechanism for heterosis in large populations. Furthermore, our future application in other species hybrid inbreeding populations will improve this method for haploid assembly using high throughput short-reads.

The Mendelian laws, developed by Gregor Mendel based on seven characteristics of the pea, are fundamental to the field of genetics. According to Mendel’s laws, diploid organisms transmit two alleles of a gene on the homologous chromosomes at an equal ratio to their offspring. Previous studies indicated, although not universal, non-Mendelian inheritance is present in crops [70], fruit flies [71], mammals [72] and birds [73]. Many factors including the meiotic drive of single chromosomes in mice, selfish genes in *Drosophila melanogaster*, or three loci (*Ga1*, *Ga2* and *Tcb1*) of unilateral cross-incompatibility (UCI) in maize [70] might cause non-Mendelian inheritance. However, the origin and mechanisms of non-Mendelian inheritance remain unclear [74]. In this study, using 12 Muscovy duck VG-haploid genomes, we evaluated segregation ratio of parental alleles from different loci with the same genotype at the genome-wide scale (Fig 7A-B). We found 38 windows totally containing 1428 genotyped loci and covering 3.18 Mb deviated significantly from the expected 1:1 segregation ratio. Further analyses indicated non-Mendelian regions had lower gene density, GC content and repeat sequence frequency, but significantly enriched DNA motifs bound by the forkhead family of transcription factors FOXC1, FOXP1 and FOXD3. Among them, *FOXC1* expression was negatively correlated with DNA repeat sequences of telomeres, which induced chromatin decompaction and increases chromatin accessibility [75]. FOXP1 increased cell cycle exit during late neurogenesis and altered chromatin accessibility [76]. While FOXD3 controlled heterochromatin-mediated repression of repeat elements and 2-cell state transcription, and promoted homologous recombination repair and genomic stability [77, 78]. Since four types of DNA motifs bound by the forkhead family of transcription factors were enriched in homologous regions of non-Mendelian regions from five divergent avian genomes (Chicken, Turkey, Mallard, Pekin duck and Kiwi; S25-26 Figs), we hypothesized these enriched DNA motifs bound by three members of the forkhead family of transcription factors might induce relatively compacted chromatin organization and low chromatin accessibility in non-Mendelian regions (Fig 7D). It might, in turn, affect allele segregation during meiosis to result in non-Mendelian regions, possibly in different birds. Previous studies had established that chromatin compactness might act as a barrier to genome editing tools, hindering the binding of CRISPR/Cas9 complexes [79]. Therefore, the heterochromatin-like features of these non-Mendelian regions might not only affect meiotic segregation but also pose challenges for precise gene editing in poultry, potentially leading to lower editing efficiency or increased off-target risks due to the inaccessibility of target loci [80]. Future studies on the origin and mechanisms of our detected non-Mendelian regions and their effect on gene editing might be informative to improve gene editing efficiency in poultry.

Hybrid phenotypic traits are determined by expression and regulation of parental genomes under specific environment [81]. For diploid organisms, two parental haploid genomes are completely homologous in homogametic individuals, but differentiate sex chromosomes in heterogametic individuals. Two sex chromosome systems have evolved in vertebrates, where the male-heterogametic XY system is for mammals and the female-heterogametic ZW system is for birds [82]. In mammals, most genes show biallelic expression, where paternal and maternal alleles are simultaneously transcribed and therefore equally contributed to the gene expression profile. In contrast, a few groups of genes show monoallelic expression, where allele-specific transcription was determined by the parent-of-origin [83–85]. Two major patterns including the X chromosome inactivation (XCI) and genomic imprinting, have been widely reported to contribute to monoallelic expression in mammals. XCI is a notable monoallelic gene expression pattern for dosage compensation in mammals, which is mediated by the noncoding *Xist* RNA coating the X chromosome and subsequently excluding RNA polymerase II from the *Xist* chromatin territory [86]. And genomic imprinting influences gene expression pattern of parent-of-origin as an epigenetic mechanism [87]. In contrast, studies on gene expression and regulatory patterns of avian parental haploid genomes are relatively limited and lack integrated multi-omics data from the same samples [10, 88]. We here integrated gene expression, chromatin conformation, chromatin accessibility and DNAm analyses to explore gene expression and regulatory patterns of parental haploid genomes in Muscovy duck. We found that paternal haploid genomes generally had lower levels of gene expression, slightly compacted chromatin conformation and lower levels of chromatin accessibility, but showed similar methylation profiles, when compared to the maternal haploid genomes (Figs 3-5). These observations suggested that, in general, Muscovy duck preferred to regulate parental allele expression through changing chromatin conformation and accessibility instead of DNA methylation at these four stages. Surprisingly, we identified a large proportion (27.24-30.13%) of genes showed monoallelic expression and 5.19% of genes were expressed in monoallelic patterns in four tissues. This observation, together with 79 SNPs being listed as potentially imprinted in chicken [88], gives rise to the question about re-evaluating genomic imprinting in birds systematically in the future.

## Conclusions

In this study, we generated high-quality chromosome-level diploid and haploid genomes for the Muscovy duck and developed an efficient, cost-effective graph-based strategy for assembling haplotype-resolved genomes. Our multi-omics analysis revealed distinct regulatory landscapes for parental alleles in which maternal haploid genomes generally exhibited more open chromatin organization and higher gene expression on autosomes. Conversely, the ZW sex-determination system displayed a unique dosage compensation mechanism where the paternal Z chromosome showed greater chromatin accessibility and transcriptional activity than the maternal Z chromosome. Furthermore, we identified specific non-Mendelian regions characterized by compacted chromatin structures and enrichment of transcription factor binding motifs, offering new insights into the constraints of allele segregation. Collectively, these findings provide a comprehensive resource for avian genomics and establish a theoretical foundation for understanding the molecular mechanisms of heterosis that is critical for optimizing poultry breeding programs.

## Abbreviations

ASE: Allele-specific expression
ATAC-seq: Assay for transposase-accessible chromatin with high-throughput sequencing
Bi-ATACPs: Biallelic ATAC-seq peaks
Bi-DNAms: Biallelic DNA methylation genes/CpG islands
BUSCO: Benchmarking Universal Single-Copy Orthologs
DeMA: Deterministic monoallelic
DNAm: DNA methylation
FDR: False discovery rate
GO: Gene Ontology
GWAS: Genome-wide association studies
Hi-C: High-throughput chromosome conformation capture
KEGG: Kyoto Encyclopedia of Genes and Genomes
Ms-ATACPs: Maternal allele-specific ATAC-seq peaks
Ms-DNAms: Maternal allele-specific DNA methylation genes/CpG islands NGS: Next-generation sequencing
ONT: Oxford Nanopore Technologies PE: Paired-end
Ps-ATACPs: Paternal allele-specific ATAC-seq peaks
Ps-DNAms: Paternal allele-specific DNA methylation genes/CpG islands
QTL: Quantitative trait loci
RaMA: Random autosomal monoallelic
RID: Relative interaction difference
SNP: Single-nucleotide polymorphism
TAD: Topologically associated domain
TSS: Transcription start site
VCF: Variant call format
VG: Variation graph
XCI: X-chromosome inactivation
Pat: Paternal
Mat: Maternal

## Declarations

### Ethics approval and consent to participate

All animal research protocols were approved by the Beijing Association for Science and Technology (approval ID SYXK, Beijing, 2007–0023). The study was performed in strict compliance with the Beijing Laboratory Animal Welfare and Ethics guidelines issued by the Beijing Administration Committee of Laboratory Animals and the Institutional Animal Care and Use Committee guidelines of China Agricultural University (CAU) (ID: SKLAB-B-2010-003).

### Availability of data and materials

The data underlying this article are available in the NCBI database under BioProject accession numbers PRJNA984447, PRJNA1019115, and PRJNA984943. The genome assembly generated in this study is available at NCBI under the name *Cairina moschata* genome assembly ASM4831997v1. The custom scripts used for analysis are available on Figshare at https://figshare.com/s/c40736465392c9db20a5.

### Consent for publication

Not applicable.

## Acknowledgments

We are grateful to the High-Performance Computing Platform of China Agricultural University for providing computational resources.

## Authors’ contributions

Y.H. conceived and supervised the project. T.L. and Y.W. performed the genome assembly, quality assessment, gene prediction, annotation, and all major bioinformatics analyses (including RNA-seq, Hi-C, ATAC-seq, and non-Mendelian region analysis). Z.Z., C.C., N.Z., J.W. (Jun Wang), M.N., J.W. (Jian Wang), and H.A. were responsible for animal care, sample collection, and experimental data generation. T.L. and Y.W. wrote the manuscript. Y.H. and H.A. revised the manuscript. All authors read and approved the final manuscript.

## Supporting information

**S1 Fig. Estimation of genome size.**

The genome size of an F1 female hybrid (Female-1) from a French Crimo Muscovy duck (male) and Yongchun Muscovy duck (female) was estimated to be 1.27 Gb based on the k-mer method.

**S2 Fig. Expanded/contracted gene families in *Cairina moschata*.**

Analysis of expanded and contracted gene families in *Cairina moschata* compared to other avian species.

**S3 Fig. Genealogy of the six offspring individuals.**

Pedigree chart showing the relationships between the six haploid genomic individuals and their parents.

**S4 Fig. BUSCO assessments of diploid and haploid genomes.**

Comparison of assembly completeness for diploid and haploid genomes at both genome and proteome levels.

**S5 Fig. BlobPlot evaluation of VG haploid genome assemblies.**

Phasing completeness evaluation of the 12 VG-haploid genomes for three full-sibling pairs (female 1-3 and male 1-3). The bubble plots illustrate the separation of paternal (red) and maternal (blue) k-mers, confirming high-quality haplotype phasing and synteny with the diploid reference.

**S6 Fig. Functional enrichment analysis.**

KEGG functional enrichment analysis of paternal and maternal allele-specific expression in varying tissues.

**S7 Fig. Hi-C resolution of individuals.**

Evaluation of Hi-C data resolution for individual samples used in chromatin architecture analysis.

**S8 Fig. Overall Hi-C resolution after merging.**

Assessment of overall Hi-C map resolution after merging biological replicates.

**S9 Fig. Hi-C A/B compartment length (Mb).**

Distribution of A/B compartment lengths across different samples and haplotypes.

**S10 Fig. Hi-C A/B compartment number.**

Comparison of the total number of A/B compartments identified in each haplotype.

**S11 Fig. Hi-C A/B compartment proportion.**

Proportional differences of A/B compartments between paternal and maternal genomes.

**S12 Fig. Functional enrichment of A/B switched regions.**

KEGG and GO enrichment analysis for regions that switch compartment status between haplotypes. (A) KEGG analysis. (B) GO analysis.

**S13 Fig. Functional enrichment of A/B variable regions.**

KEGG and GO analysis for regions showing variable compartment strength. (A) KEGG analysis. (B) GO analysis.

**S14 Fig. TAD size per individual.**

Comparison of Topologically Associating Domain (TAD) sizes across individuals and haplotypes.

**S15 Fig. Enrichment analysis of Ps-ATACPs and Ms-ATACPs.**

Functional enrichment of genes associated with paternal-specific (Ps) and maternal-specific (Ms) ATAC-seq peaks.

**S16 Fig. Ratio of male to female FPKM values on the Z chromosome.**

The scatter plots display the log2-transformed ratios of male-to-female gene expression (FPKM) for Z-linked genes across different tissues (brain, liver, spleen, and muscle) and developmental stages (Weeks 1, 2, 4, and 7). The distribution of these ratios illustrates the degree of dosage compensation between sexes.

**S17 Fig. Z chromosome interaction scaling.**

Intra-chromosomal Hi-C interaction scaling plots (log-log) comparing paternal, maternal, and female Z chromosomes. (A) Paternal-Z vs. maternal-Z. (B) Female-Z vs. maternal-Z. (C) Female-Z vs. paternal-Z.

**S18 Fig. Z chromosome compartment changes.**

Analysis of A/B compartment switching between paternal and maternal Z haplotypes.

**S19 Fig. Functional analysis of Z-linked A/B compartments.**

KEGG and GO annotation of genes located in A/B switched regions on the Z chromosome. (A) KEGG analysis. (B) GO analysis.

**S20 Fig. Statistical charts of haploid Z-chromosome TADs.**

Comparison of TAD numbers, length distribution, and boundary strength on the haploid Z chromosome. (A) Comparison of TAD numbers on the haploid Z chromosome. (B) Length distribution of haploid Z-chromosome TADs. (C) Boundary strength comparison of haploid Z-chromosome TADs.

**S21 Fig. Haploid Z-chromosome ATAC-seq distribution map.**

Differential heatmap of ATAC-seq peaks between paternal and maternal alleles in male individuals.

**S22 Fig. Identification of genes with allelically biased TSS peak clustering.**

Analysis identifying genes with exclusive paternal or maternal peak clustering on the Z chromosome. (A–B) KEGG pathway enrichment analysis of genes associated with paternal-specific (A) and maternal-specific (B) ATAC-seq peaks. (C–D) Gene Ontology (GO) enrichment analysis of genes associated with paternal-specific (C) and maternal-specific (D) ATAC-seq peaks

**S23 Fig. Sliding window segmentation of the whole genome.**

The curve illustrates the decay of statistical significance (*P*-value) for non-Mendelian detection as window size decreases from 10 Mb to 100 kb.

**S24 Fig. Genotype distribution map of the whole genome.**

Genotypes were categorized into Mendelian and non-Mendelian regions based on the analysis of six haploid individuals and their parents.

**S25 Fig. Non-Mendelian regions in different species.**

Characteristics of non-Mendelian regions across different species. (A-E) Scatter plots showing the frequency of repeated sequences in Muscovy duck compared to Pekin duck, Mallard, Turkey, Chicken and Kiwi.

**S26 Fig. Syntenic analysis of motif enrichment.**

Syntenic analyses demonstrate that the above four types of DNA motifs were also enriched in homologous regions of non-Mendelian regions from five divergent avian genomes (Chicken, Turkey, Mallard, Pekin duck, and Kiwi).

S1 Table. Statistics of normal and ultra-long Nanopore reads.

S2 Table. Statistics of contig lengths assembled with Nanopore reads.

S3 Table. Statistics of clean Illumina genomic data.

S4 Table. Statistics of Nanopore assembly contig lengths after polishing with Illumina data.

S5 Table. Statistics of Bionano BNX reads.

S6 Table. Statistics of scaffolds generated from Bionano and Nanopore reads.

S7 Table. Statistics of Hi-C reads.

S8 Table. Chromosome lengths of the Muscovy duck (*Cairina moschata*, SKLFABB.Cai Mos.1.0) and Pekin duck (SKLA2.0) genomes.

S9 Table. Statistics of the TrioCanu haploid genomes.

S10 Table. RNA-Seq sequencing statistics.

S11 Table. Statistics of the transcripts.

S12 Table. Expanded/contracted gene families in *Cairina moschata*.

S13 Table. Species used in the maximum likelihood tree.

S14 Table. Statistics of the VG haploid genomes.

S15 Table. Merqury statistics of the haploid genomes.

S16 Table. Differences between the VG haploid genomes and TrioCanu haploid genomes.

S17 Table. BUSCO score statistics for quality.

S18 Table. Hi-C data statistics for quality.

S19 Table. Statistics of ATAC-seq peaks on the Z chromosome.

S20 Table. A/B compartment statistics on the Z chromosome.

S21 Table. TAD statistics on the Z chromosome.

S22 Table. ATAC-seq peak statistics on the Z chromosome.

S23 Table. Statistics of motif frequency.

