## supporting file for "Exploring genetic, expression and regulatory patterns of parental alleles in Muscovy duck (*Cairina moschata*) using haplotype-resolved assemblies"

**S Tables**

**S1 Table** Statistics of standard and ultra-long Nanopore reads

| Index | Libraries | |
| --- | --- | --- |
|  | Normal | Ultra-long1 |
| Total Reads Base (Gb) | 123 | 17.19 |
| Clean Reads Base (Gb) | 106 | 14.02 |
| Clean Read Count (Millions) | 3.93 | 0.29 |
| Clean Reads Mean Length (kb) | 25.18 | 58.91 |
| Clean Reads N50 Length (kb) | 32.66 | 90.72 |
| Clean Reads Max Length (kb) | 236.66 | 578.1 |

**S2 Table** Statistics of contig lengths assembled with Nanopore reads

| Index | Length (Mb) | Number |
| --- | --- | --- |
| N10 | 149.93 | 1 |
| N20 | 80.42 | 2 |
| N30 | 71.55 | 4 |
| N40 | 42.68 | 10 |
| N50 | 39.41 | 18 |
| N60 | 26.37 | 21 |
| N70 | 22.21 | 27 |
| N80 | 10.67 | 30 |
| N90 | 6.53 | 37 |
| Min. | 0.3 | # |
| Max. | 113.41 | # |
| Ave. | 5.82 | # |
| Total | 1181.96 | 203 |

**S3 Table** Statistics of clean Illumina genomic data

| Paired-end libraries (bp) | Paired-end insert size (bp) | Average Read length (bp) | Number of reads (×10^6^) | Total data (Gb) | Sequence coverage (fold) |
| --- | --- | --- | --- | --- | --- |
| 400 | 400 | 150 | 891.16 | 117.13 | 100 |

**S4 Table** Statistics of Nanopore assembly contig lengths after polishing with Illumina data

| Index | Length (Mb) | Number |
| --- | --- | --- |
| N10 | 149.99 | 1 |
| N20 | 81.42 | 2 |
| N30 | 71.85 | 4 |
| N40 | 42.98 | 10 |
| N50 | 40.41 | 18 |
| N60 | 27.37 | 21 |
| N70 | 22.95 | 27 |
| N80 | 11.47 | 30 |
| N90 | 6.91 | 37 |
| Min. | 0.9 | # |
| Max. | 117.41 | # |
| Ave. | 5.88 | # |
| Total | 1184.61 | 203 |

**S5 Table** Statistics of Bionano BNX reads

| Index | Size |
| --- | --- |
| Maps | 470793 |
| Sites | 23048393 |
| Length (kb) | 1138907998 |
| Average length (kb) | 242.961 |
| Label Density (each 100 kb) | 15.731 |
| N50 (Mb) | 0.2461 |

**S6 Table** Statistics of scaffolds generated from Bionano and Nanopore reads

| Index | Scaffold length (Mb) |
| --- | --- |
| Number of scaffolds | 154 |
| Min length | 0.06 |
| Median length | 5.15 |
| Mean length | 8.01 |
| N50 length | 85.82 |
| Max length | 205.77 |
| Total length | 1231.17 |

Note: The unit for length is Mb..

**S7 Table** Statistics of Hi-C reads

| Total Reads  Base (Gb) | Total Reads Number (Mb) | Clean Reads  Base (Gb) | Clean Reads Number (Mb) | Clean Reads  Rate (%) |
| --- | --- | --- | --- | --- |
| 289.13 | 1942.81 | 274.19 | 1777.65 | 94.80 |

**S8 Table** Chromosome lengths of the Muscovy duck (*Cairina moschata*, SKLFABB.CaiMos.1.0) and Pekin duck (SKLA2.0) genomes

| Chromosome | Size (Mb) | GC% | Number of coding genes | SKLA2.0 |
| --- | --- | --- | --- | --- |
| chr1 | 209.37 | 39.75 | 2200 | 207.84 |
| chr2 | 157.85 | 39.17 | 1371 | 160.52 |
| chr3 | 121.77 | 39.29 | 1366 | 118.87 |
| chr4 | 76.99 | 39.78 | 806 | 76.57 |
| chr5 | 66.86 | 40.55 | 989 | 66.4 |
| chr6 | 37.68 | 41.5 | 543 | 38.14 |
| chr7 | 42.15 | 38.48 | 506 | 39.93 |
| chr8 | 34.77 | 40.34 | 524 | 32.71 |
| chr9 | 29.36 | 39.1 | 455 | 26.7 |
| chr10 | 23.06 | 39.43 | 351 | 22.33 |
| chr11 | 22.29 | 43.38 | 415 | 22.15 |
| chr12 | 23.52 | 39.53 | 376 | 21.81 |
| chr13 | 22.24 | 43.3 | 365 | 22.48 |
| chr14 | 22.44 | 40.84 | 378 | 20.47 |
| chr15 | 20.16 | 39.94 | 425 | 18.17 |
| chr16 | 16.6 | 45.81 | 381 | 15.6 |
| chr17 | 4.38 | 55.09 | 99 | 4.13 |
| chr18 | 11.99 | 48.09 | 307 | 12.03 |
| chr19 | 15.31 | 41.09 | 326 | 12.62 |
| chr20 | 7.94 | 47.11 | 262 | 11.97 |
| chr21 | 16.14 | 45.66 | 371 | 16.13 |
| chr22 | 9.27 | 45.45 | 264 | 8.53 |
| chr23 | 6.45 | 48.79 | 202 | 5.9 |
| chr24 | 7.62 | 50.43 | 250 | 8.03 |
| chr25 | 10.32 | 37.67 | 185 | 7.89 |
| chr26 | 3.16 | 57.27 | 324 | 3.24 |
| chr27 | 9.61 | 45.67 | 360 | 6.83 |
| chr28 | 6.64 | 51.68 | 343 | 7.25 |
| chr29 | 6.38 | 50.23 | 314 | 5.93 |
| chr30 | 3.31 | 57.17 | 218 | 5.46 |
| chr31 | 1.73 | 47.66 | 18 | 1.95 |
| chr32 | 3.2 | 45 | 131 | 5.24 |
| chr33 | 4.25 | 27.23 | 61 | 2.2 |
| chr34 | 3.55 | 36.4 | 24 | 4.82 |
| chr35 | 7.2 | 49.84 | 76 | 3.26 |
| chr36 | 3.44 | 32.04 | 185 | 3.57 |
| chr37 | 7.78 | 49.3 | 142 | 2.45 |
| chr38 | 8.55 | 30.34 | 24 | 1.91 |
| chr39 | 5.96 | 52.29 | 30 | 1.79 |
| chrW | 19.13 | 42.51 | 100 | # |
| chrZ | 85.79 | 40.06 | 781 | 85.48 |

**S9 Table** Statistics of the TrioCanu haploid genomes

|  | paternal | | maternal | |
| --- | --- | --- | --- | --- |
|  | contig | chromosome | contig | chromosome |
| length | 1,115,688,179 | 1,105,373,511 | 1,080,983,596 | 1,060,805,918 |
| number | 254 | 137 | 298 | 199 |
| min | 121,380 | 227 | 61,532 | 210 |
| max | 47,152,608 | 201,065,607 | 32,253,600 | 202,328,177 |
| N50 | 11,512,528 | 81,825,781 | 10,524,097 | 75,400,788 |
| avg | 4,392,473.10 | 8,068,419.80 | 3,627,461.70 | 5,330,683 |

**S10 Table** RNA-Seq sequencing statistics

| GEO code | Sample | | | Mapping ratio (%) |
| --- | --- | --- | --- | --- |
| SRR25592441 | | female | 87.98 | |
| SRR25592440 | | female | 91.77 | |
| SRR25592316 | | female | 92.63 | |
| SRR25592352 | | female | 79.23 | |
| SRR25592313 | | female | 90.52 | |
| SRR25592314 | | female | 90.01 | |
| SRR25592373 | | female | 96.8 | |
| SRR25592398 | | male | 92.01 | |
| SRR25592432 | | male | 83.89 | |
| SRR25592423 | | male | 89.33 | |
| SRR25592436 | | male | 88.59 | |
| SRR25592427 | | male | 90.26 | |
| SRR25592322 | | female | 92.45 | |
| SRR25592388 | | male | 80.69 | |
| SRR25592372 | | female | 92.05 | |
| SRR25592422 | | male | 86.39 | |
| SRR25592357 | | female | 88.62 | |
| SRR25592299 | | female | 91.6 | |
| SRR25592334 | | female | 81.94 | |
| SRR25592356 | | female | 90.91 | |
| SRR25592301 | | female | 88.42 | |
| SRR25592335 | | female | 89.5 | |
| SRR25592300 | | female | 93.31 | |
| SRR25592426 | | male | 83.34 | |
| SRR25592406 | | female | 89.43 | |
| SRR25592411 | | male | 88.66 | |
| SRR25592349 | | female | 87.43 | |
| SRR25592429 | | male | 91.9 | |
| SRR25592359 | | female | 83.26 | |
| SRR25592307 | | female | 88.73 | |
| SRR25592327 | | female | 86.96 | |
| SRR25592386 | | male | 87.08 | |
| SRR25592323 | | female | 93.13 | |
| SRR25592303 | | female | 83.38 | |
| SRR25592400 | | male | 90.63 | |
| SRR25592347 | | female | 88.99 | |
| SRR25592439 | | female | 88.28 | |
| SRR25592366 | | female | 92.4 | |
| SRR25592339 | | female | 84.16 | |
| SRR25592328 | | female | 89.02 | |
| SRR25592393 | | male | 88.65 | |
| SRR25592311 | | female | 87.21 | |
| SRR25592391 | | male | 92.74 | |
| SRR25592421 | | male | 84.12 | |
| SRR25592424 | | male | 92.09 | |
| SRR25592306 | | female | 89.37 | |
| SRR25592295 | | male | 87.3 | |
| SRR25592410 | | male | 92.76 | |
| SRR25592404 | | male | 83.44 | |
| SRR25592367 | | female | 88.12 | |
| SRR25592408 | | male | 90.38 | |
| SRR25592302 | | female | 79.21 | |
| SRR25592370 | | female | 93.39 | |
| SRR25592361 | | female | 88.13 | |
| SRR25592333 | | female | 88.67 | |
| SRR25592309 | | female | 88.41 | |
| SRR25592379 | | female | 88.46 | |
| SRR25592395 | | male | 92.04 | |
| SRR25592430 | | male | 86.23 | |
| SRR25592324 | | female | 89.65 | |
| SRR25592390 | | male | 91.75 | |
| SRR25592351 | | female | 89.42 | |
| SRR25592402 | | male | 94.43 | |
| SRR25592344 | | female | 88.74 | |
| SRR25592376 | | female | 92.54 | |
| SRR25592428 | | female | 88.26 | |
| SRR25592317 | | female | 87.06 | |
| SRR25592298 | | female | 91.66 | |
| SRR25592354 | | female | 82.23 | |
| SRR25592394 | | female | 93.01 | |
| SRR25592355 | | female | 87.15 | |
| SRR25592365 | | female | 90.72 | |
| SRR25592381 | | female | 92.68 | |
| SRR25592315 | | female | 84.22 | |
| SRR25592368 | | female | 91.41 | |
| SRR25592369 | | female | 88.26 | |
| SRR25592371 | | female | 88.95 | |
| SRR25592337 | | female | 91.68 | |
| SRR25592340 | | female | 86.41 | |
| SRR25592413 | | male | 92.12 | |
| SRR25592431 | | male | 89.18 | |
| SRR25592380 | | female | 87.12 | |
| SRR25592304 | | female | 93.21 | |
| SRR25592417 | | female | 84.4 | |
| SRR25592319 | | female | 91.53 | |
| SRR25592419 | | male | 88.48 | |
| SRR25592416 | | male | 88.15 | |
| SRR25592296 | | male | 93.54 | |
| SRR25592397 | | male | 81.2 | |
| SRR25592346 | | female | 84.79 | |
| SRR25592345 | | female | 88.45 | |
| SRR25592433 | | male | 87.71 | |
| SRR25592435 | | male | 91.68 | |
| SRR25592437 | | male | 80.43 | |
| SRR25592329 | | female | 73.75 | |
| SRR25592364 | | female | 87.94 | |
| SRR25592348 | | female | 89.67 | |
| SRR25592392 | | male | 93.12 | |
| SRR25592332 | | female | 86.18 | |
| SRR25592401 | | male | 93.32 | |
| SRR25592378 | | female | 88.73 | |
| SRR25592377 | | female | 95.84 | |
| SRR25592409 | | male | 93.01 | |
| SRR25592403 | | male | 85.63 | |
| SRR25592414 | | male | 91.65 | |
| SRR25592420 | | male | 88.6 | |
| SRR25592375 | | female | 85.86 | |
| SRR25592396 | | male | 92.56 | |
| SRR25592387 | | male | 87 | |
| SRR25592330 | | female | 89.34 | |
| SRR25592343 | | female | 89.71 | |
| SRR25592341 | | female | 96.48 | |
| SRR25592412 | | male | 92.02 | |
| SRR25592425 | | male | 88.13 | |
| SRR25592326 | | female | 91.22 | |
| SRR25592318 | | female | 89.21 | |
| SRR25592407 | | male | 89.89 | |
| SRR25592353 | | female | 92.78 | |
| SRR25592438 | | male | 86.16 | |
| SRR25592342 | | female | 93.5 | |
| SRR25592338 | | female | 89.32 | |
| SRR25592415 | | male | 85.84 | |
| SRR25592405 | | male | 91.33 | |
| SRR25592363 | | female | 85.92 | |
| SRR25592383 | | female | 91.59 | |
| SRR25592389 | | male | 88.09 | |
| SRR25592350 | | female | 82.23 | |
| SRR25592308 | | female | 91.52 | |
| SRR25592360 | | female | 80.48 | |
| SRR25592362 | | female | 92.32 | |
| SRR25592320 | | female | 88.64 | |
| SRR25592374 | | female | 88.23 | |
| SRR25592310 | | female | 88 | |
| SRR25592297 | | female | 82.71 | |
| SRR25592385 | | female | 91.2 | |
| SRR25592434 | | male | 89.15 | |
| SRR25592325 | | female | 85.29 | |
| SRR25592418 | | male | 91.94 | |
| SRR25592399 | | male | 77.21 | |
| SRR25592336 | | female | 93.17 | |
| SRR25592382 | | female | 88.34 | |
| SRR25592312 | | female | 83.99 | |
| SRR25592305 | | female | 91.81 | |
| SRR25592331 | | female | 81.74 | |
| SRR25592321 | | female | 86.09 | |
| SRR25592294 | | male | 84.64 | |
| SRR25592358 | | female | 87.66 | |
| SRR25592384 | | female | 91.53 | |

**S11 Table** Statistics of the transcripts

| Genome | Gene count | Transcript count |
| --- | --- | --- |
| SKLFABB.CaiMos1.0 | 16893 | 46141 |
| TrioCanu-paternal | 16613 | 43139 |
| TrioCanu-maternal | 16571 | 43524 |
| female-1-maternal | 16521 | 43194 |
| female-1-paternal | 16318 | 42746 |
| female-2-maternal | 16531 | 43798 |
| female-2-paternal | 16388 | 42467 |
| female-3-maternal | 16513 | 43746 |
| female-3-paternal | 16314 | 43414 |
| female-4-maternal | 16532 | 43831 |
| female-4-paternal | 16519 | 43791 |
| female-5-maternal | 16522 | 43770 |
| female-5-paternal | 16547 | 43718 |
| female-6-maternal | 16513 | 43801 |
| female-6-paternal | 16519 | 43798 |

**S12 Table** Expanded/contracted gene families in *Cairina moschata*

| ID | Family name | Number of genes | | | | |
| --- | --- | --- | --- | --- | --- | --- |
|  |  | Anas platyrhynchos | Cairina moschata | Pekin duck | Muscovy ducks and duck common ancestor | P-value (* 0 denotes P < 1×10⁻²⁰⁰) |
| OG0015459 | 1-acyl-sn-glycerol-3-phosphate acyltransferase alpha | 2 | 1 | 2 | 2 | 0 |
| OG0015462 | 40S ribosomal protein | 2 | 1 | 2 | 2 | 0 |
| OG0000040 | Very low-density lipoprotein receptor | 10 | 1 | 16 | 9 | 0 |
| OG0001563 | Death-inducer obliterator | 8 | 1 | 9 | 7 | 0 |
| group58 | Serine/threonine-protein phosphatase | 8 | 1 | 8 | 7 | 0 |
| group37 | Heat shock transcription factor, Y-linked | 7 | 1 | 7 | 6 | 0.01 |
| group12 | GTPase-activating Rap/Ran-GAP domain-like protein | 6 | 1 | 6 | 5 | 0 |
| OG0000099 | Cohesin subunit | 4 | 1 | 4 | 4 | 0 |
| group02 | Interferon alpha-inducible protein 27-like protein | 4 | 1 | 4 | 4 | 0 |
| OG0014605 | Putative uncharacterized protein C6orf52 | 4 | 1 | 5 | 4 | 0 |
| OG0000313 | Zinc finger | 23 | 6 | 29 | 21 | 0 |
| PTHR45710 | Coiled-coil domain-containing protein | 12 | 3 | 12 | 10 | 0 |
| OG0000651 | Collagen alpha-1(XI) chain | 3 | 1 | 3 | 3 | 0 |
| group42 | HLA class II histocompatibility antigen | 3 | 1 | 4 | 3 | 0 |
| PTHR46746 | Killer cell lectin-like receptor subfamily B member | 3 | 1 | 5 | 3 | 0 |
| group39 | Mitochondrial import receptor subunit TOM5 homolog | 3 | 1 | 5 | 3 | 0 |
| PTHR24247 | Mucin-5AC | 3 | 1 | 4 | 3 | 0 |
| PTHR24253 | Potassium/sodium hyperpolarization-activated cyclic nucleotide-gated channel | 3 | 1 | 4 | 3 | 0 |
| OG0001920 | Protocadherin gamma | 3 | 1 | 3 | 3 | 0.021 |
| OG0000224 | SLA class II histocompatibility antigen | 6 | 2 | 7 | 6 | 0 |
| OG0000080 | TRPM8 channel-associated factor | 3 | 1 | 3 | 3 | 0.04 |
| OG0000017 | Ubiquitin carboxyl-terminal hydrolase protein | 3 | 1 | 3 | 3 | 0.02 |
| PTHR24064 | GTPase IMAP family member | 11 | 4 | 20 | 11 | 0 |
| OG0000015 | ATPase family AAA domain-containing protein | 15 | 5 | 16 | 13 | 0.005 |
| PTHR24399 | Ankyrin repeat domain-containing protein | 5 | 2 | 13 | 5 | 0.007 |
| group01 | Outer dense fiber protein | 6 | 2 | 6 | 5 | 0 |
| OG0000709 | Antigen WC1.1 | 14 | 6 | 18 | 14 | 0 |
| PTHR45615 | C-C motif chemokine 3-like protein | 8 | 3 | 8 | 7 | 0.001 |
| group36 | Kinesin-like protein | 8 | 3 | 9 | 7 | 0 |
| group06 | Maestro heat-like repeat-containing protein family member | 13 | 6 | 19 | 13 | 0 |
| OG0000025 | Beta-crystallin | 1 | 1 | 3 | 2 | 0 |
| OG0001322 | Chitinase-3-like protein | 2 | 1 | 3 | 2 | 0 |
| OG0000081 | Embryonic protein | 2 | 1 | 2 | 2 | 0 |
| OG0000933 | Epidermal differentiation-specific protein | 4 | 2 | 5 | 4 | 0 |
| OG0000018 | Gag-pol polyprotein | 4 | 2 | 4 | 4 | 0 |
| group38 | Guanylate-binding protein | 2 | 2 | 8 | 4 | 0 |
| group14 | IgGFc-binding protein | 1 | 1 | 2 | 2 | 0 |
| group03 | Interferon-induced protein with tetratricopeptide repeats | 2 | 1 | 2 | 2 | 0 |
| group47 | Low affinity immunoglobulin gamma Fc region receptor | 2 | 1 | 2 | 2 | 0 |
| group08 | Natural killer cell receptor | 2 | 1 | 7 | 2 | 0.01 |
| group43 | Neuronal acetylcholine receptor | 2 | 1 | 2 | 2 | 0 |
| group45 | Nuclear pore complex protein Nup153 | 2 | 1 | 10 | 2 | 0 |
| group48 | Phthioceranic/hydroxyphthioceranic acid synthase | 2 | 1 | 3 | 2 | 0 |
| group49 | Pleckstrin homology domain-containing family H member | 2 | 1 | 2 | 2 | 0 |
| OG0001182 | Putative exonuclease GOR | 2 | 1 | 2 | 2 | 0 |
| OG0000188 | Small EDRK-rich factor | 2 | 1 | 2 | 2 | 0 |
| OG0000960 | SUN domain-containing protein | 2 | 1 | 2 | 2 | 0 |
| OG0000229 | Taste receptor type 2 member | 4 | 2 | 4 | 4 | 0 |
| OG0000036 | von Willebrand factor A domain-containing protein | 1 | 1 | 6 | 2 | 0 |
| PTHR24118 | Deleted in malignant brain tumors 1 protein | 26 | 14 | 34 | 24 | 0 |
| OG0001456 | C-type lectin domain family | 10 | 6 | 14 | 10 | 0 |
| group41 | NAD(P)(+)--arginine ADP-ribosyltransferase | 10 | 6 | 17 | 10 | 0 |
| OG0000496 | Cytochrome P450 | 3 | 2 | 3 | 3 | 0.004 |
| group15 | Heterogeneous nuclear ribonucleoprotein A0 | 2 | 2 | 3 | 3 | 0 |
| PTHR24100 | Nuclear GTPase SLIP-GC | 6 | 4 | 9 | 6 | 0 |
| OG0008117 | Retinol dehydrogenase | 2 | 2 | 4 | 3 | 0.011 |
| OG0000056 | T cell receptor alpha variable | 6 | 4 | 7 | 6 | 0 |
| group57 | Scale keratin | 4 | 5 | 4 | 4 | 0 |
| OG0000491 | Avidin | 6 | 7 | 6 | 5 | 0 |
| OG0000588 | Envelope glycoprotein gp95 | 5 | 6 | 5 | 5 | 0.037 |
| OG0000103 | Putative butyrophilin subfamily 2 member | 5 | 6 | 5 | 5 | 0 |
| OG0000006 | Tyrosine-protein phosphatase non-receptor type substrate | 5 | 9 | 8 | 5 | 0 |
| OG0015471 | Claw keratin | 23 | 25 | 23 | 21 | 0 |
| PTHR24271 | Histone | 19 | 21 | 20 | 20 | 0 |
| OG0000153 | E3 ubiquitin-protein ligase | 8 | 11 | 10 | 8 | 0 |
| OG0000002 | Butyrophilin subfamily 2 member | 23 | 52 | 50 | 26 | 0.03 |
| OG0000088 | Feather keratin | 64 | 77 | 73 | 62 | 0 |
| OG0001264 | Protocadherin alpha-2 | 16 | 19 | 25 | 18 | 0 |
| group46 | Olfactory receptor | 232 | 253 | 292 | 238 | 0.001 |

**S13 Table** Species used in the maximum likelihood tree

| Abbreviation | Species name | Common name |
| --- | --- | --- |
| Apl_dom | Anas platyrhynchos | Pekin duck |
| Cat | Cygnus atratus | Black swan |
| Gga | Gallus gallus | Chicken |
| Hsa | Homo sapiens | Human |
| Tgu | Taeniopygia guttata | Zebra finch |
| Apl_wild | Anas platyrhynchos | Mallard |
| Cmo | Cairina moschata | Muscovy Duck |
| Ptr | Pan troglodytes | Chimpanzee |
| Xla | Xenopus laevis | African clawed frog |
| Aca | Anolis carolinensis | Green anole |
| Mga | Meleagris gallopavo | Turkey |
| Nme | Numida meleagris | Helmeted Guineafowl |

**S14 Table** Statistics of the VG haploid genomes

| Genome | Number | Length (bp) | Scaffold N50 (bp) | Contig N50 (bp) |
| --- | --- | --- | --- | --- |
| female-1-maternal | 39 | 1100132706 | 82343468 | 10138132 |
| female-1-paternal | 39 | 1050976157 | 75911107 | 11526613 |
| female-2-maternal | 39 | 1099661200 | 82325729 | 11243364 |
| female-2-paternal | 39 | 1050684157 | 75903297 | 10331482 |
| female-3-maternal | 39 | 1099661200 | 82325729 | 12334941 |
| female-3-paternal | 39 | 1050592415 | 75892640 | 11356490 |
| female-4-maternal | 39 | 1098047164 | 81825781 | 10349754 |
| female-4-paternal | 39 | 1119566356 | 82316269 | 11364751 |
| female-5-maternal | 39 | 1099661200 | 82325729 | 11336474 |
| female-5-paternal | 39 | 1119172926 | 82295035 | 12234972 |
| female-6-maternal | 39 | 1099651200 | 82323729 | 10316481 |
| female-6-paternal | 39 | 1119266940 | 82303239 | 11036447 |

**S15 Table** Merqury statistics of the VG haploid genomes

| Sample | Block_size | Hap-mer count | Site-specific count | Gene count | K | Sequence length | Proportion |
| --- | --- | --- | --- | --- | --- | --- | --- |
| TrioCanu-Paternal | 100kb | 89942146 | 80226499 | 16891 | 23 | 933313523 | 0.78 |
| TrioCanu-Maternal | 100kb | 88132441 | 79223114 | 16566 | 23 | 925443571 | 0.77 |
| female-1-paternal | 100kb | 10141911 | 83885787 | 16898 | 23 | 899334253 | 0.75 |
| female-1-maternal | 100kb | 10031755 | 80296853 | 16512 | 23 | 900296853 | 0.75 |
| female-2-paternal | 100kb | 10131641 | 83468827 | 16544 | 23 | 887931564 | 0.74 |
| female-2-maternal | 100kb | 10020945 | 80286373 | 16466 | 23 | 886634617 | 0.74 |
| female-3-paternal | 100kb | 99864318 | 88316489 | 16781 | 23 | 901316769 | 0.75 |
| female-3-maternal | 100kb | 99791318 | 87265629 | 16472 | 23 | 897443121 | 0.75 |
| female-4-paternal | 100kb | 10101298 | 80991948 | 16766 | 23 | 916398745 | 0.76 |
| female-4-maternal | 100kb | 10018544 | 87911935 | 16513 | 23 | 915313986 | 0.76 |
| female-5-paternal | 100kb | 99874361 | 80136955 | 16810 | 23 | 894657331 | 0.75 |
| female-5-maternal | 100kb | 99786643 | 80976643 | 16523 | 23 | 886392446 | 0.74 |
| female-6-paternal | 100kb | 101148975 | 82316854 | 16846 | 23 | 899745613 | 0.75 |
| female-6-maternal | 100kb | 101156842 | 81337842 | 16500 | 23 | 894683156 | 0.75 |

**S16 Table** Sequences flanking CDS in diploid and haploid genomes

| Sample | Sequence flanking CDS (kb) | | | | | | | | | |
| --- | --- | --- | --- | --- | --- | --- | --- | --- | --- | --- |
|  | 1 | 2 | 3 | 4 | 5 | 6 | 7 | 8 | 9 | 10 |
| GRCg7B | 0.78 | 1.17 | 1.49 | 1.8 | 2.07 | 2.35 | 2.66 | 2.96 | 3.2 | 3.41 |
| bTaeGut1_v1.p | 1.6 | 2.25 | 2.82 | 3.29 | 3.74 | 4.05 | 4.48 | 4.81 | 5.17 | 5.53 |
| SLKA2.0 | 0.28 | 0.45 | 0.59 | 0.68 | 0.74 | 0.84 | 0.94 | 1.04 | 1.12 | 1.26 |
| SKLFABB.CaiMos1.0 | 0.36 | 0.49 | 0.67 | 0.73 | 0.83 | 0.86 | 0.99 | 1.06 | 1.19 | 1.3 |
| TrioCanu-Paternal | 0.52 | 0.61 | 0.71 | 0.82 | 1.03 | 1.33 | 1.59 | 1.61 | 1.84 | 2.24 |
| TrioCanu-Maternal | 0.51 | 0.66 | 0.77 | 0.81 | 1.05 | 1.37 | 1.57 | 1.64 | 1.93 | 2.27 |
| female-1-paternal | 0.47 | 0.55 | 0.68 | 0.78 | 0.9 | 1.21 | 1.52 | 1.53 | 1.89 | 2.06 |
| female-2-maternal | 0.49 | 0.58 | 0.7 | 0.81 | 0.92 | 1.31 | 1.61 | 1.63 | 1.9 | 2.12 |
| female-2-paternal | 0.48 | 0.57 | 0.71 | 0.8 | 0.91 | 1.32 | 1.61 | 1.64 | 2 | 2.1 |
| female-3-maternal | 0.48 | 0.55 | 0.7 | 0.82 | 0.91 | 1.24 | 1.58 | 1.67 | 2.21 | 2.33 |
| female-3-paternal | 0.47 | 0.56 | 0.71 | 0.83 | 0.92 | 1.28 | 1.59 | 1.69 | 2.15 | 2.31 |
| female-4-maternal | 0.49 | 0.58 | 0.74 | 0.82 | 0.92 | 1.31 | 1.6 | 1.72 | 2.1 | 2.21 |
| female-4-paternal | 0.49 | 0.59 | 0.75 | 0.81 | 0.93 | 1.29 | 1.58 | 1.71 | 2.13 | 2.28 |
| female-5-maternal | 0.48 | 0.55 | 0.78 | 0.81 | 0.94 | 1.21 | 1.57 | 1.61 | 1.91 | 2.03 |
| female-5-paternal | 0.49 | 0.54 | 0.77 | 0.8 | 0.94 | 1.19 | 1.6 | 1.63 | 1.91 | 2.12 |
| female-6-maternal | 0.48 | 0.57 | 0.67 | 0.76 | 0.89 | 1.14 | 1.48 | 1.71 | 1.9 | 2.1 |
| female-6-paternal | 0.47 | 0.56 | 0.66 | 0.79 | 0.91 | 1.2 | 1.43 | 1.7 | 1.88 | 2.13 |

**S17 Table** Differences between the VG haploid genomes and Trio-binning haploid genomes

|  | Vg-female-1-paternal | Vg-female-1-maternal | Trio-female-1-paternal | Trio-female-1-maternal |
| --- | --- | --- | --- | --- |
| Covered genes | 15458 | 15360 | 15773 | 15618 |
| Uncovered genes | 1063 | 1104 | 731 | 812 |
| Length (bp) | 1025928546 | 972615629 | 1057947624 | 991467323 |
| Total (bp) | 1099851207 | 1040684157 | 1105373511 | 1060805918 |
| Uncovered length (bp) | 73922661 | 68068528 | 47425887 | 69338595 |
| Density | 1.50673E-05 | 1.57925E-05 | 1.49091E-05 | 1.57524E-05 |
| Uncovered density | 1.43799E-05 | 1.62141E-05 | 1.54135E-05 | 1.17106E-05 |
| Size | 564 | 554 | 431 | 412 |
| Max | 1766 | 1352 | 3721 | 3964 |
| Uncovered size | 41 | 39 | 44 | 41 |
| Max | 612 | 101 | 1712 | 1314 |
| Number | 1818397 | 1756751 | 4396370 | 4192843 |

**S18 Table** BUSCO score statistics for quality/duplication

| Genome | S | D | F | M | C |
| --- | --- | --- | --- | --- | --- |
| Vg-maternal | 94 | 0.5 | 1.1 | 4.4 | 94.5 |
| Vg-paternal | 93.8 | 0.5 | 1 | 4.7 | 94.3 |
| Trio-maternal | 93.9 | 0.4 | 1.1 | 4.5 | 94.4 |
| Trio-paternal | 93.7 | 0.5 | 0.9 | 4.9 | 94.2 |
| Protein | S | D | F | M | C |
| VG-maternal | 94.1 | 0.7 | 0.4 | 4.8 | 94.8 |
| VG-paternal | 93.9 | 0.7 | 0.3 | 5.1 | 94.6 |
| Trio-maternal | 94 | 0.7 | 0.4 | 4.9 | 94.7 |
| Trio-paternal | 93.8 | 0.7 | 0.4 | 5.1 | 94.5 |

**S19 Table** Hi-C data statistics for quality/duplication

| Sample | Valid pairs | Unique valid pairs | Unique (%) | Cis/Trans |
| --- | --- | --- | --- | --- |
| Female-1 | 554,176,912 | 363,096,713 | 65.52 | 5.01 |
| Female-2 | 561,829,953 | 312,208,905 | 55.57 | 4.22 |
| Female-3 | 534,291,842 | 401,894,324 | 75.22 | 4.06 |
| Male-1 | 571,926,641 | 386,336,446 | 67.55 | 4.15 |
| Male-2 | 505,531,792 | 387,287,906 | 76.61 | 4.42 |
| Male-3 | 516,697,183 | 320,558,932 | 62.04 | 4.66 |

Note: This table shows filtering statistics for quality control and duplicate removal of Hi-C data.

**S20 Table** Statistics of ATAC-seq peaks on the Z chromosome

| Sample | Number | Gene count | TSS(+-500bp) |
| --- | --- | --- | --- |
| female-1-paternal | 54566 | 15189 | 14683 |
| female-1-maternal | 54312 | 15087 | 14389 |
| female-2-paternal | 51511 | 14257 | 12464 |
| female-2-maternal | 54489 | 14346 | 12296 |
| female-3-paternal | 60973 | 15653 | 13852 |
| female-3-maternal | 65750 | 15542 | 13583 |
| male-1-paternal | 59211 | 15212 | 13046 |
| male-1-maternal | 59221 | 14590 | 13138 |
| male-2-paternal | 63508 | 14318 | 12139 |
| male-2-maternal | 64919 | 14564 | 12703 |
| male-3-paternal | 51147 | 14595 | 13839 |
| male-3-maternal | 52355 | 14780 | 13579 |
| Merge-paternal | 59563 | 13922 | 13007 |
| Merge-maternal | 59266 | 14073 | 12788 |

**S21 Table** A/B compartment statistics on the Z chromosome

| Sample | Number | Size | Gene |
| --- | --- | --- | --- |
| male-paternal-A | 543 | 30764794 | 603 |
| male-paternal-B | 313 | 55585205 | 158 |
| male-maternal-A | 295 | 56628079 | 158 |
| male-maternal-B | 533 | 32641968 | 603 |
| female-paternal-A | 533 | 55585205 | 603 |
| female-paternal-B | 331 | 34519142 | 158 |

**S22 Table** TAD statistics on the Z chromosome

| Sample | TAD number | Average size (bp) | Gene count |
| --- | --- | --- | --- |
| male-paternal | 123 | 81420000 | 762 |
| male-maternal | 123 | 81560000 | 760 |
| female-paternal | 123 | 81320000 | 762 |

**S23 Table** ATAC-seq peak statistics on the Z chromosome

| sample | number | size | gene |
| --- | --- | --- | --- |
| male-active | 3566 | 82324864 | 713 |
| male-inactive | 3505 | 82302407 | 702 |
| female-paternal | 3572 | 81320000 | 711 |

**S24 Table** Statistics of motif frequency

| Sample | Area | AAACAAACAAACAAA | ATTAAAAAAAAAAAA | AGAAAAAAAAAAAAA | TAAATAAATAAATAA |
| --- | --- | --- | --- | --- | --- |
|  |  | FOXL1 | FOXL1 | FOXL1 | FOXL1 |
| Muscovy duck | Non-Mendelian | 0.70044 | 0.65052 | 0.63492 | 0.70668 |
|  | Mendelian | 0.60996 | 0.49452 | 0.48516 | 0.624 |
| Pekin duck | Non-Mendelian | 0.69108 | 0.6552 | 0.62712 | 0.70356 |
|  | Mendelian | 0.62244 | 0.50076 | 0.47268 | 0.62556 |
| Mallard | Non-Mendelian | 0.69732 | 0.65676 | 0.63648 | 0.7098 |
|  | Mendelian | 0.6084 | 0.49296 | 0.4914 | 0.63024 |
| Turkey | Non-Mendelian | 0.6552 | 0.62088 | 0.62556 | 0.68796 |
|  | Mendelian | 0.59436 | 0.51792 | 0.50076 | 0.62868 |
| Kiwi | Non-Mendelian | 0.6474 | 0.64116 | 0.64272 | 0.68952 |
|  | Mendelian | 0.61152 | 0.50232 | 0.49608 | 0.57096 |
| Chicken | Non-Mendelian | 0.62556 | 0.62556 | 0.59436 | 0.68796 |
|  | Mendelian | 0.60528 | 0.49608 | 0.48984 | 0.64116 |

**S Figures**
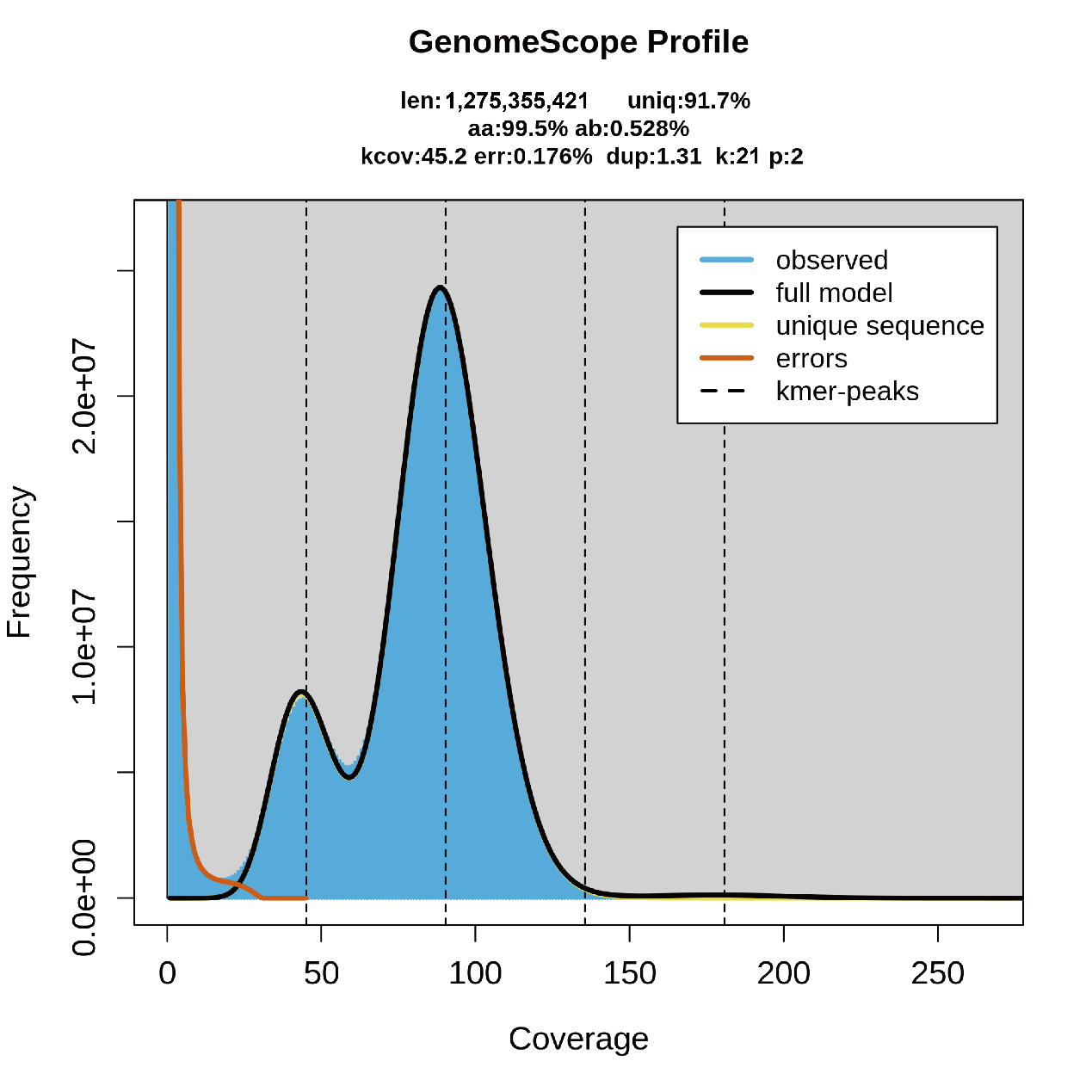

**S1 Fig. Estimation of genome size.** The genome size of an F1 female hybrid (Female-1) from a French Crimo Muscovy duck (male) and Yongchun Muscovy duck (female) was estimated to be 1.32 Gb based on the k-mer method.

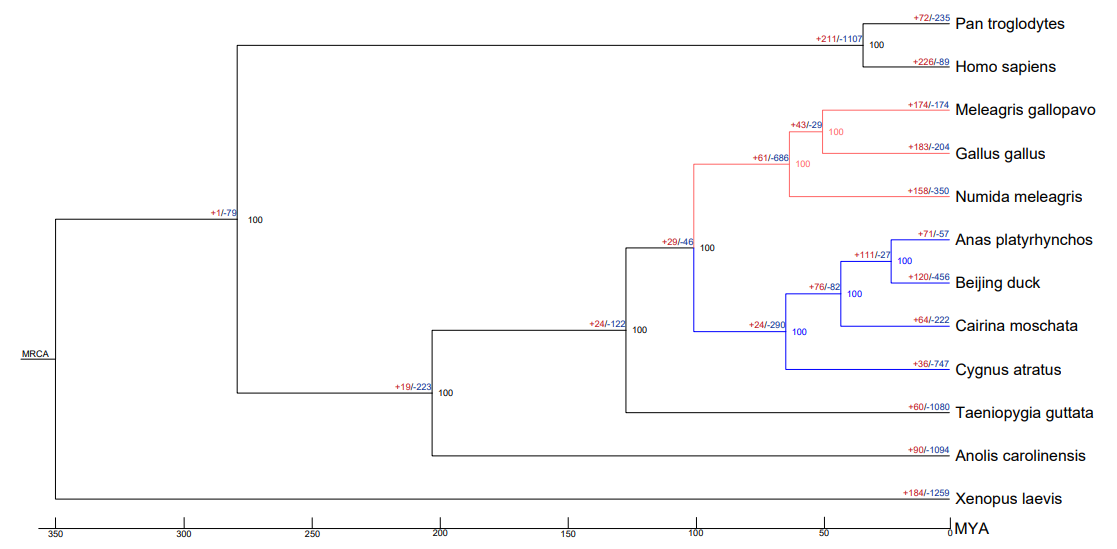
**S2 Fig. Expanded/contracted gene families in *Cairina moschata*.**

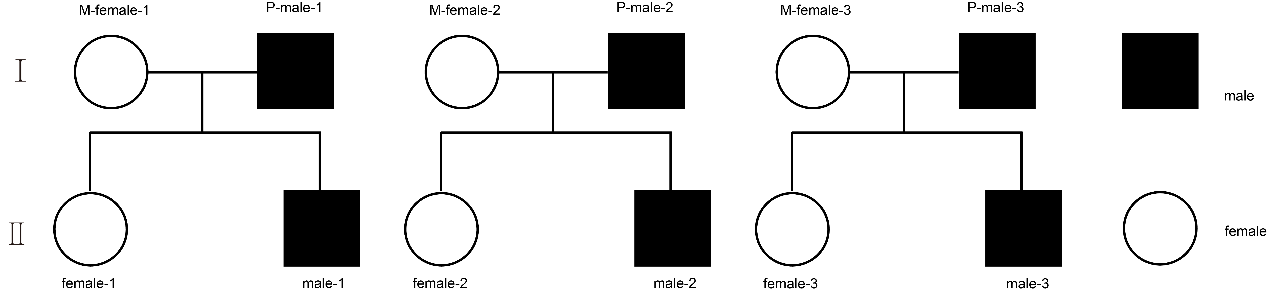

**S3 Fig. Genealogy of the six offspring individuals.** Genealogical relationships between six haploid genomic individuals and their parents.

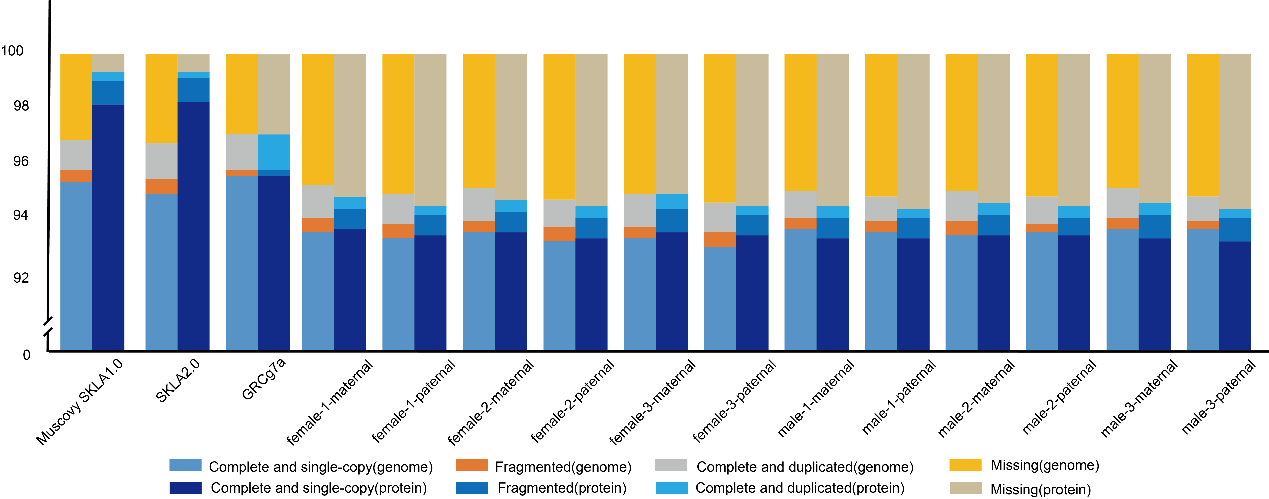

**S4 Fig.** **BUSCO assessments of diploid and haploid genomes (genome and proteome).**

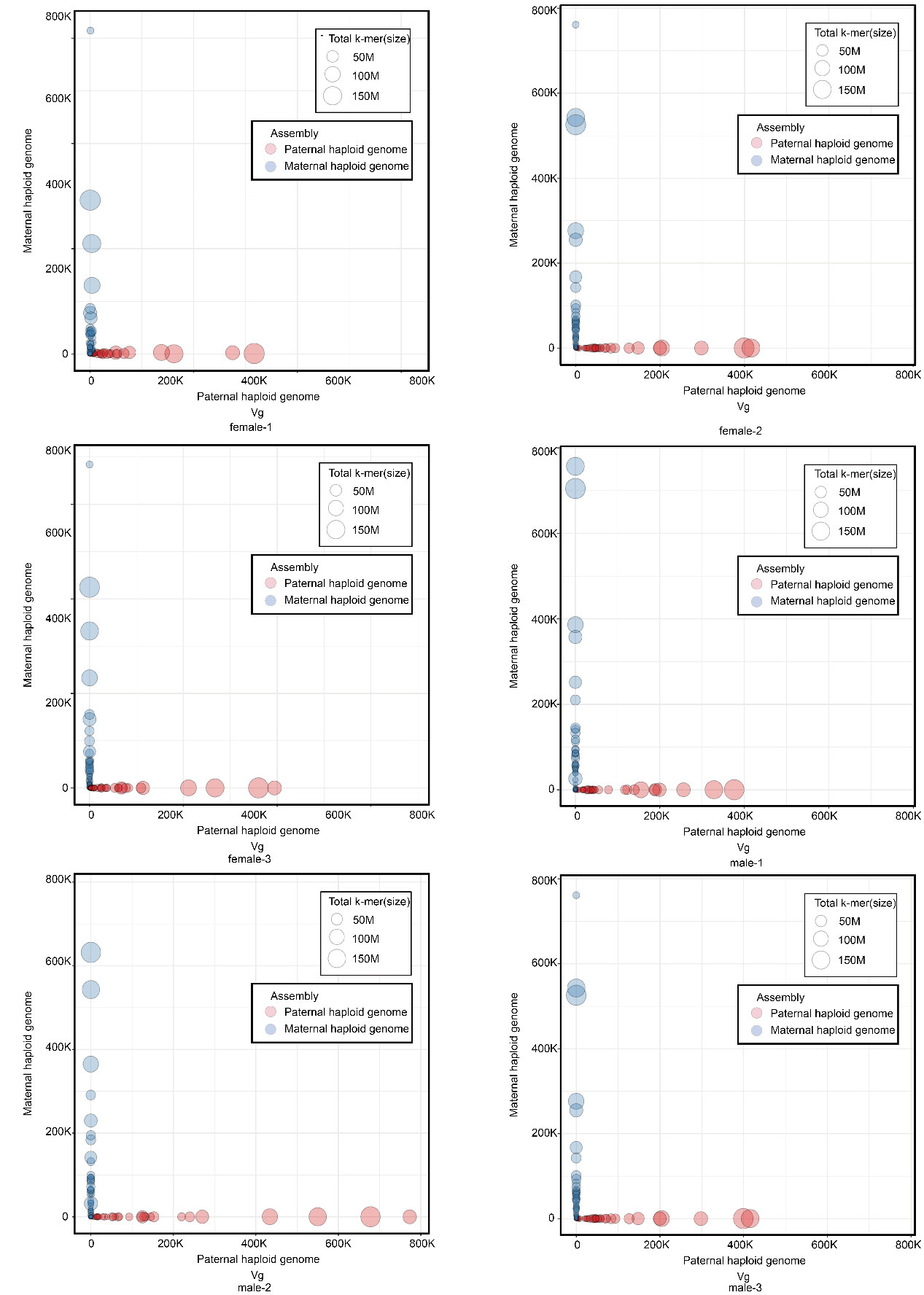

**S5 Fig. BlobPlot evaluation of VG haploid genome assemblies.**

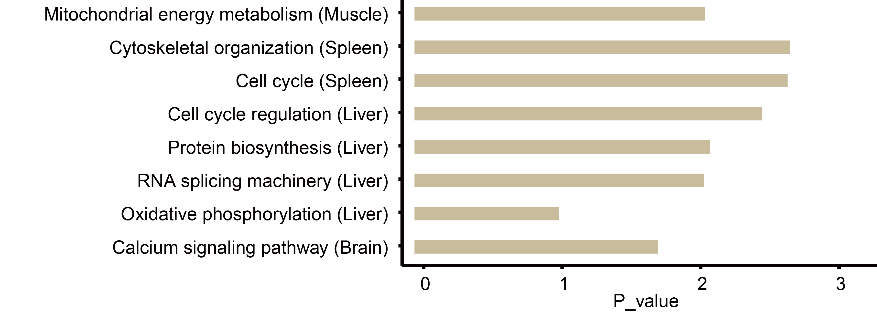

**S6 Fig. Functional enrichment analysis.** KEGG functional enrichment analysis of paternal and maternal allele-specific expression.

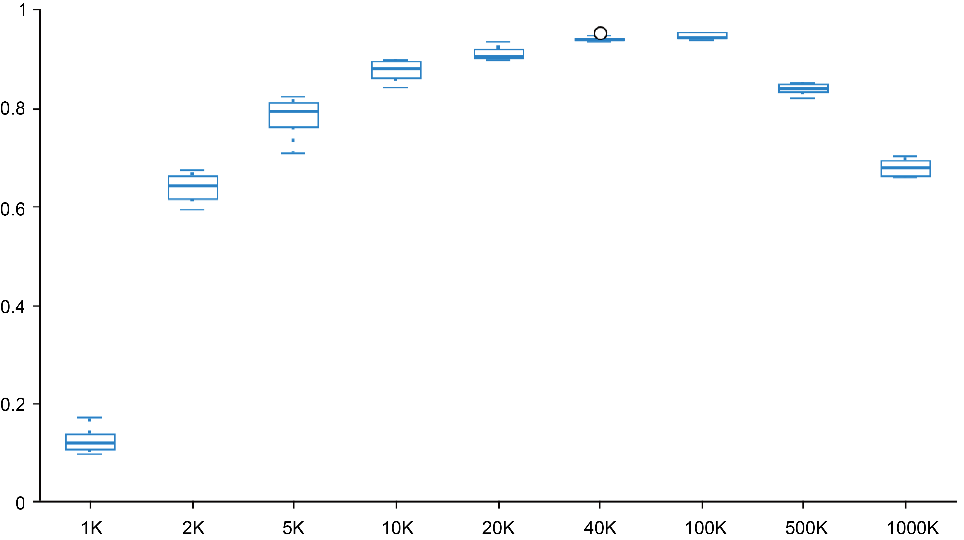

**S7 Fig.** **Hi-C resolution of individuals.**

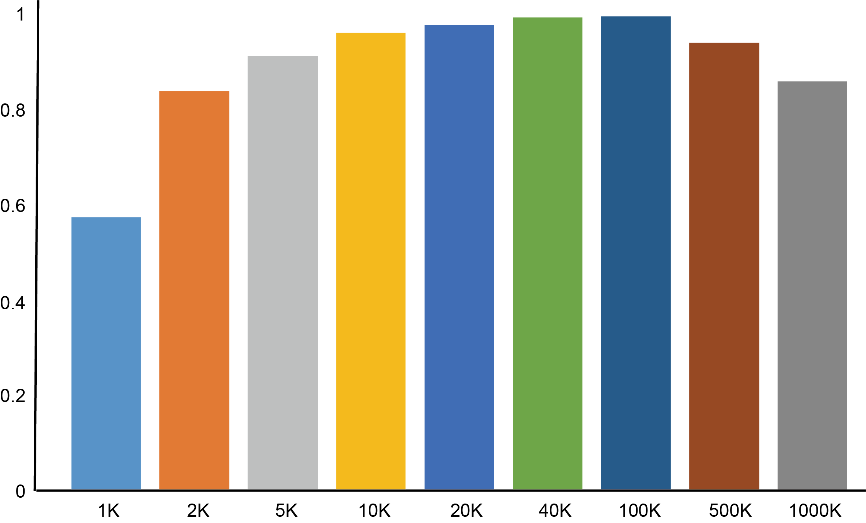

**S8 Fig.** **Overall Hi-C resolution after merging.**

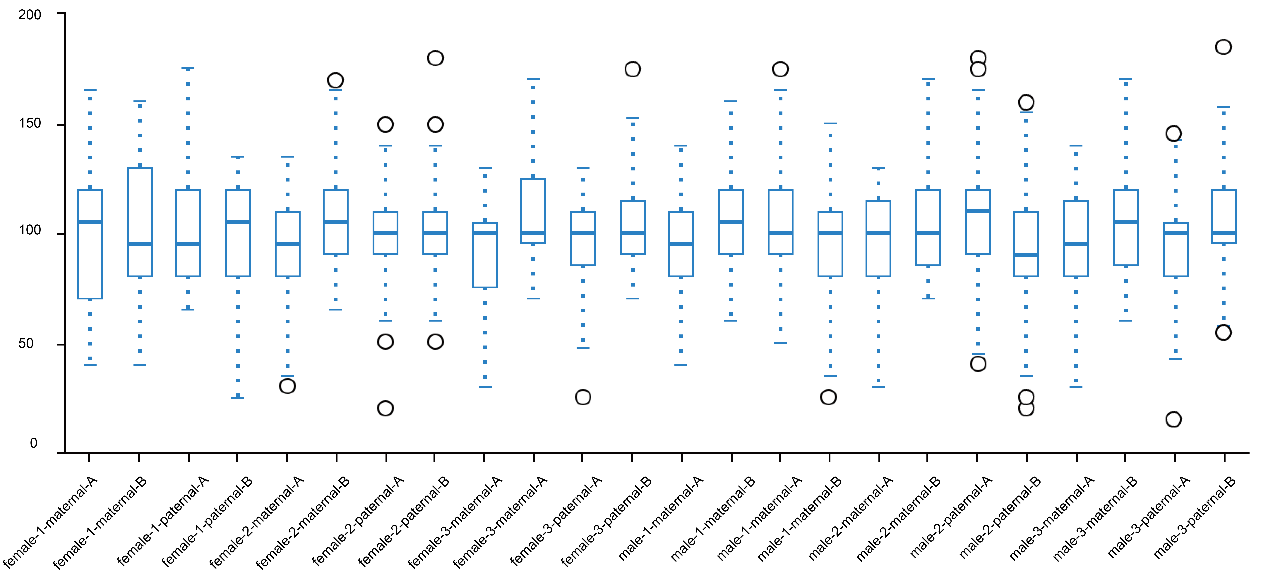
 **S9 Fig. Hi-C A/B compartment length (Mb)**.

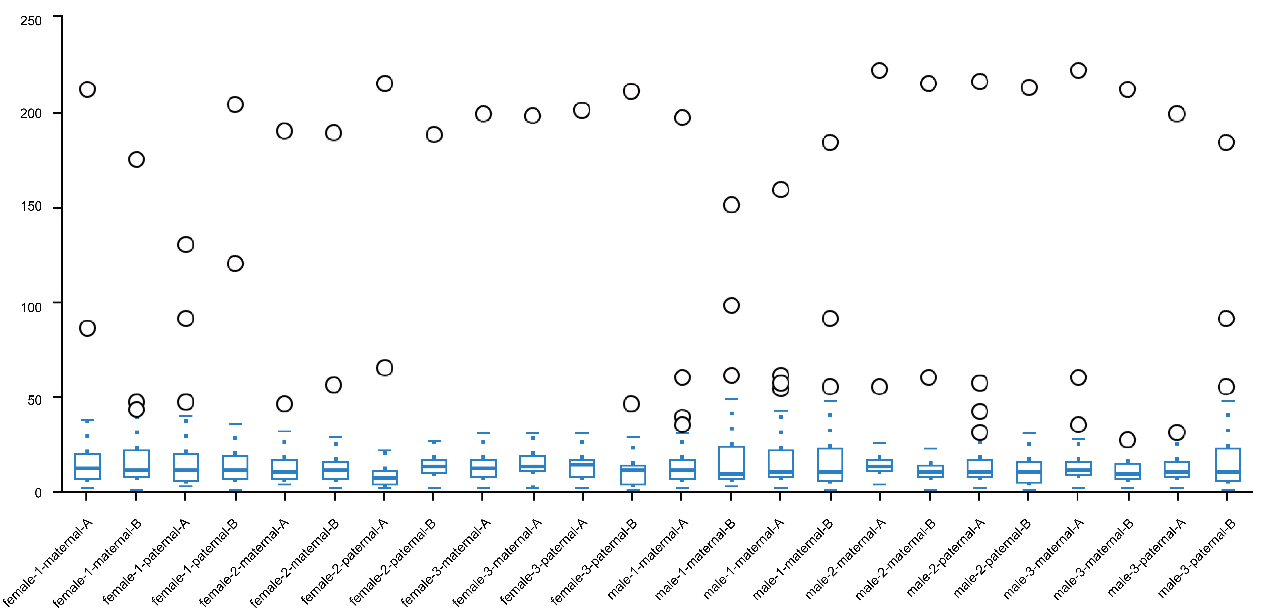

**S10 Fig. Hi-C A/B compartment number.**

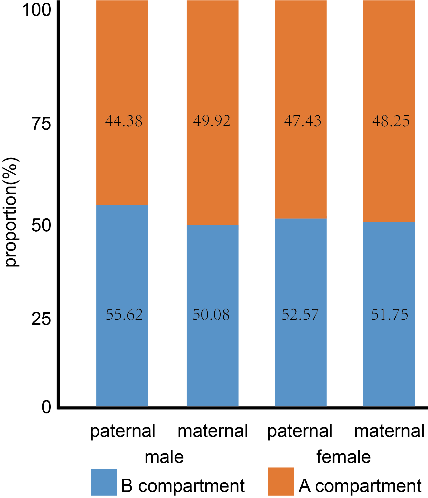

**S11 Fig. Hi-C A/B compartment proportion.** Proportional differences of A/B compartments between paternal and maternal genomes.

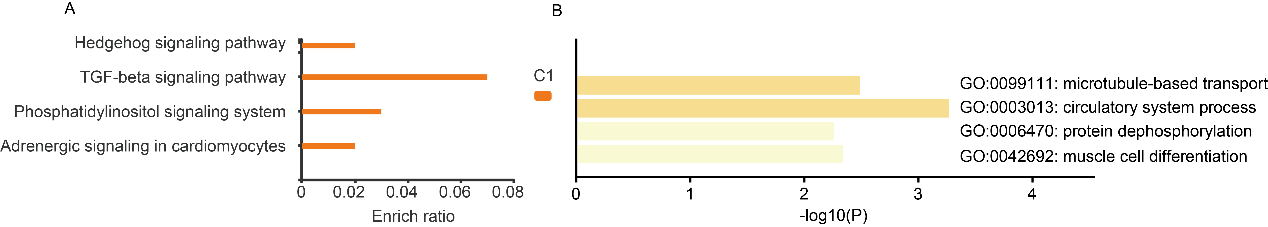

**S12 Fig. KEGG and GO functional enrichment analysis of A/B switched regions.** (A) KEGG analysis. (B) GO analysis.

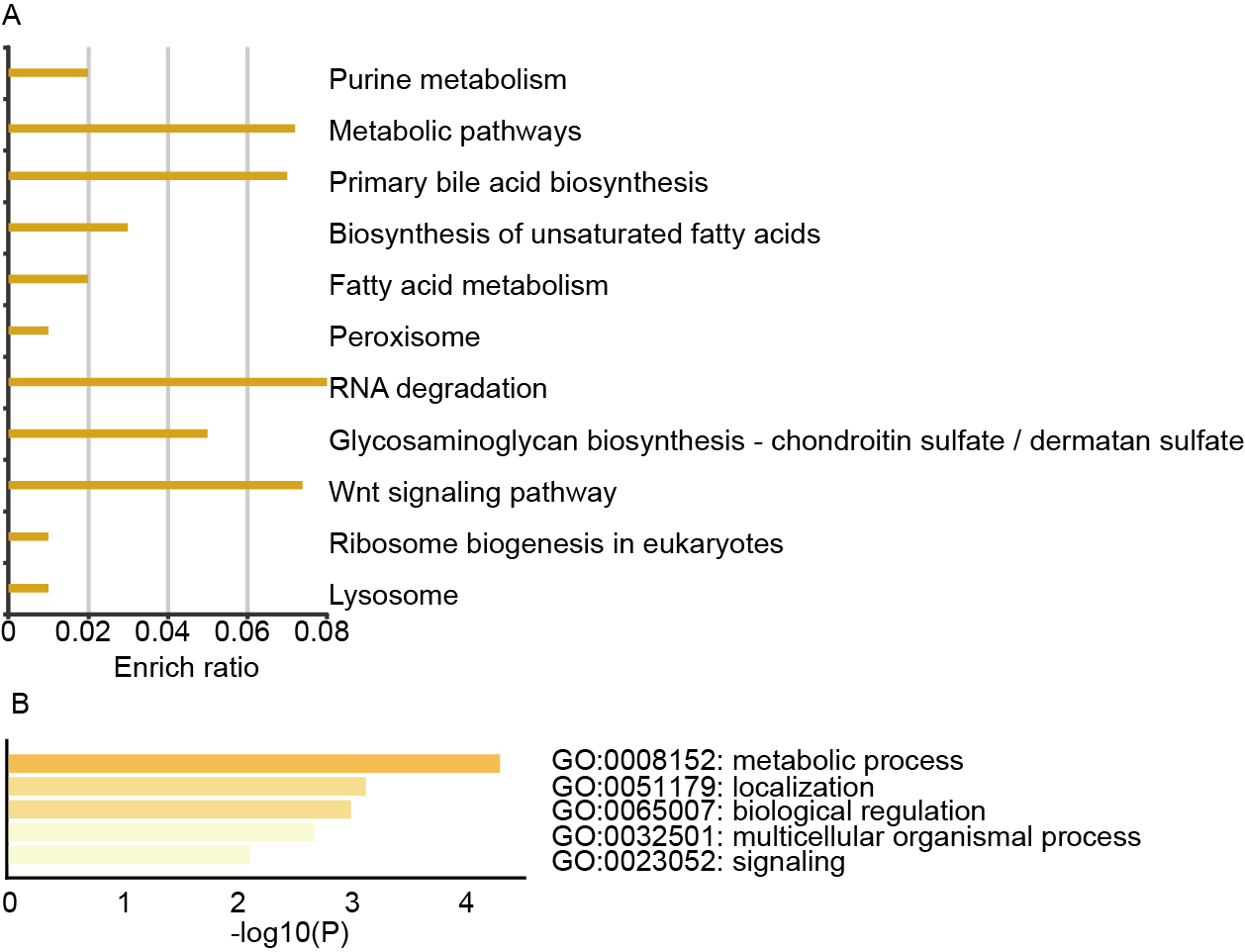
**S13 Fig. KEGG and GO functional enrichment analysis of A/B variable regions.** (A) KEGG analysis. (B) GO analysis.

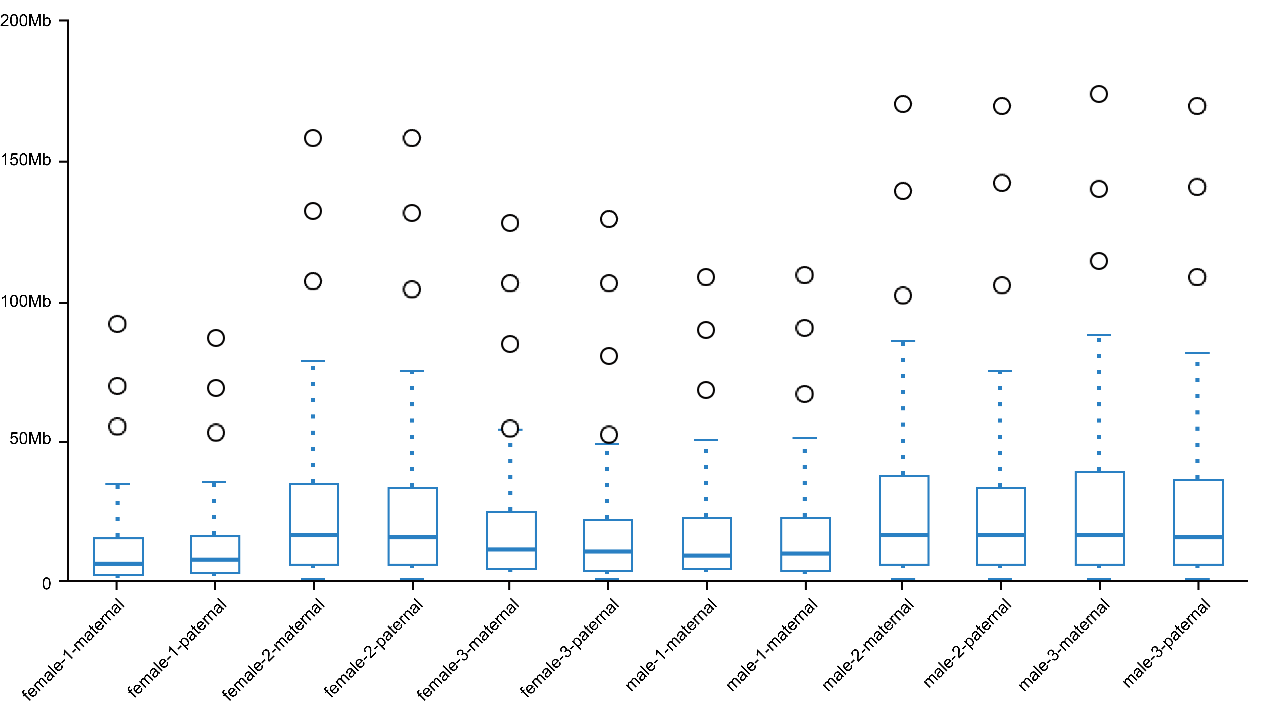

**S14 Fig. TAD size per individual.**

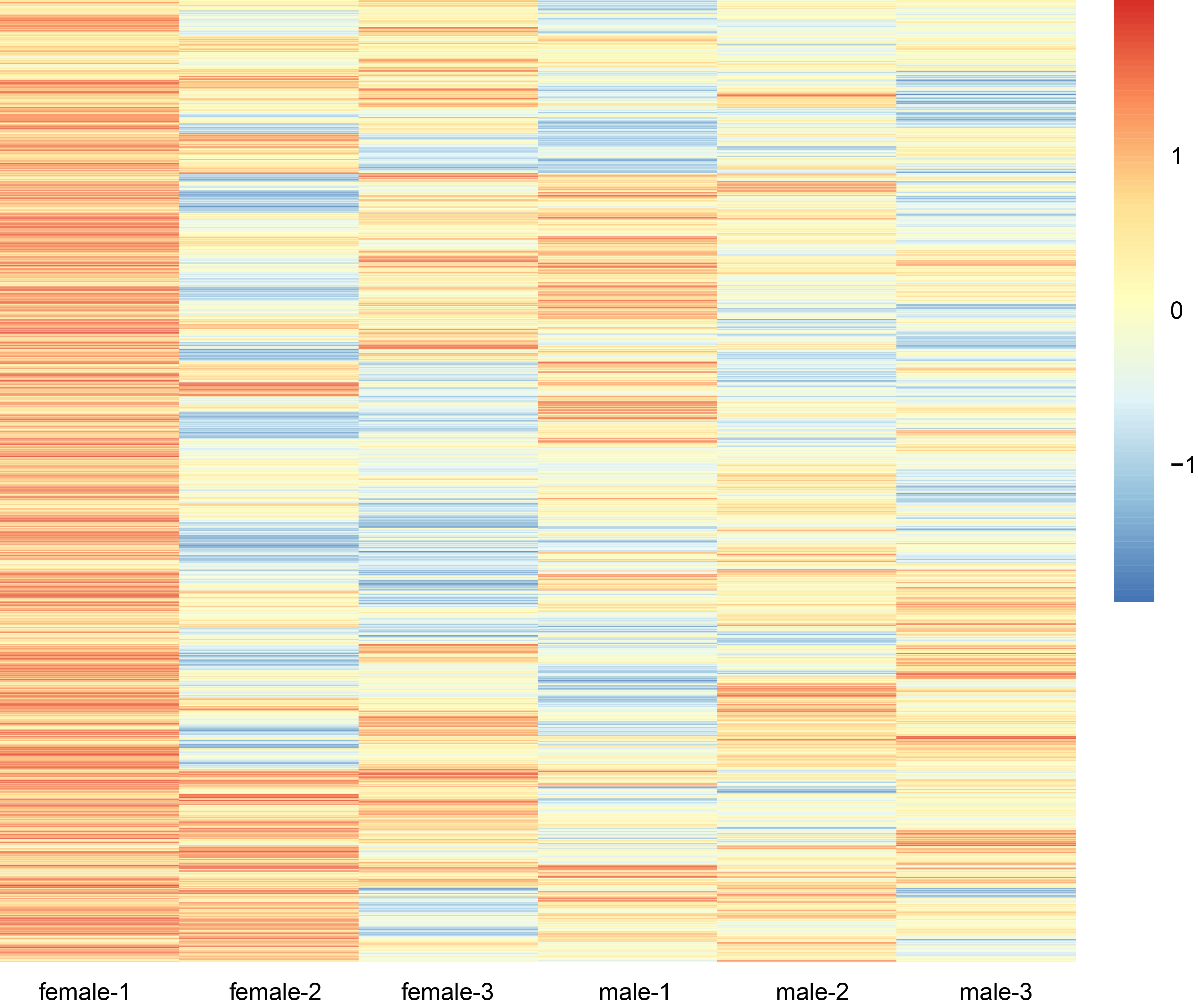

**S15 Fig. Haploid whole-genome ATAC-seq distribution map.** Heatmap of differences between individual paternal and maternal ATAC-seq peaks.

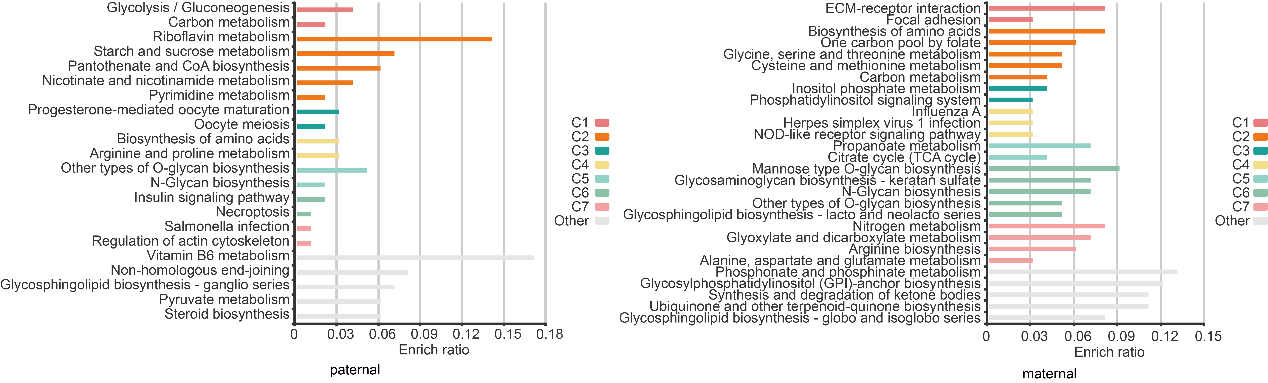

**S16 Fig. KEGG and GO functional enrichment analysis of Ps-ATACPs and Ms-ATACPs.**

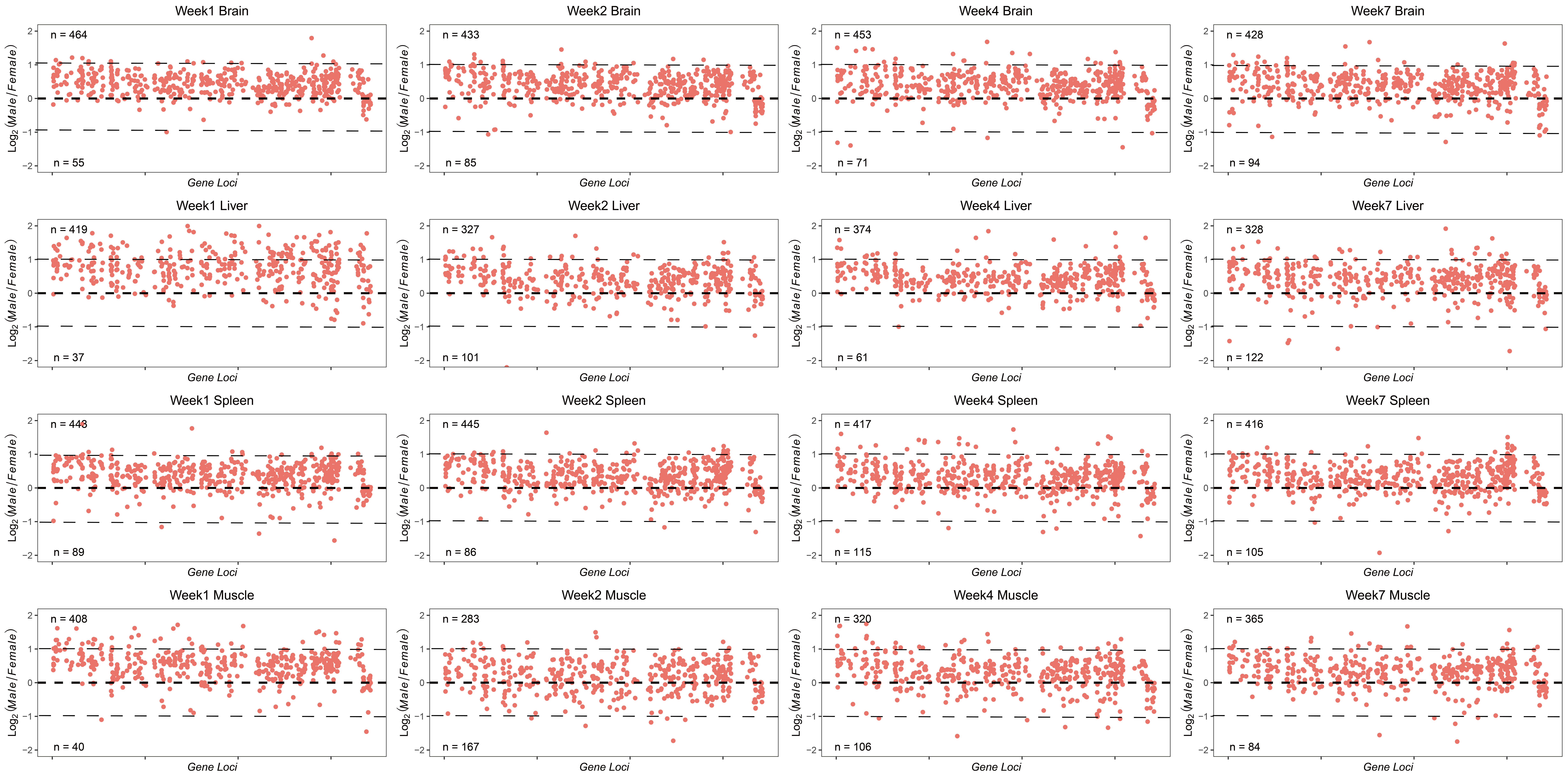

**S17 Fig. Ratio of male to female FPKM values on the Z chromosome.** The scatter plots display the log2-transformed ratios of male-to-female gene expression (FPKM) for Z-linked genes across different tissues (brain, liver, spleen, and muscle) and developmental stages (Weeks 1, 2, 4, and 7). The distribution of these ratios illustrates the degree of dosage compensation between sexes.

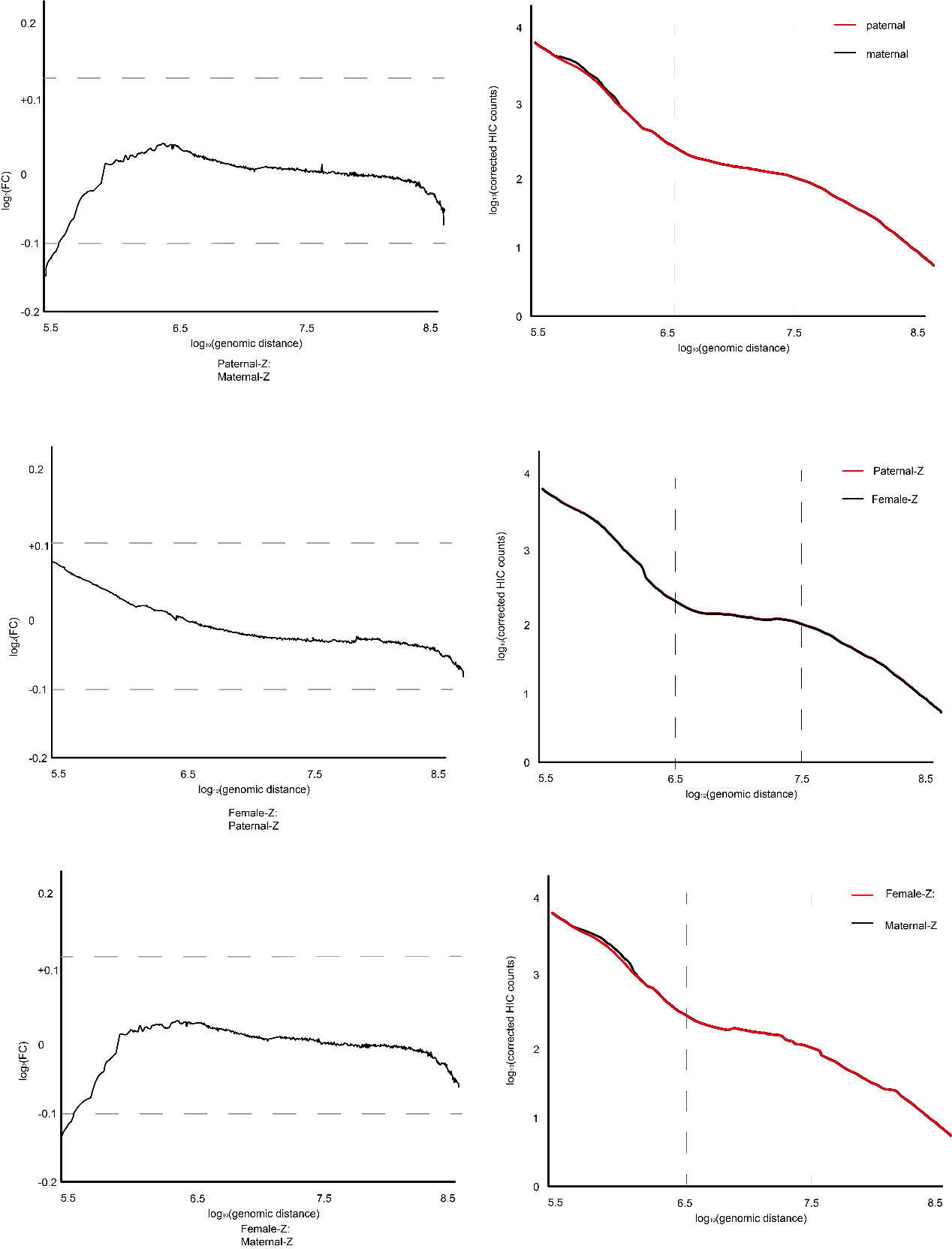

**S18 Fig. Intra-chromosomal Hi-C interaction scaling plots (log-log) comparing the paternal, maternal, and female Z chromosomes.**

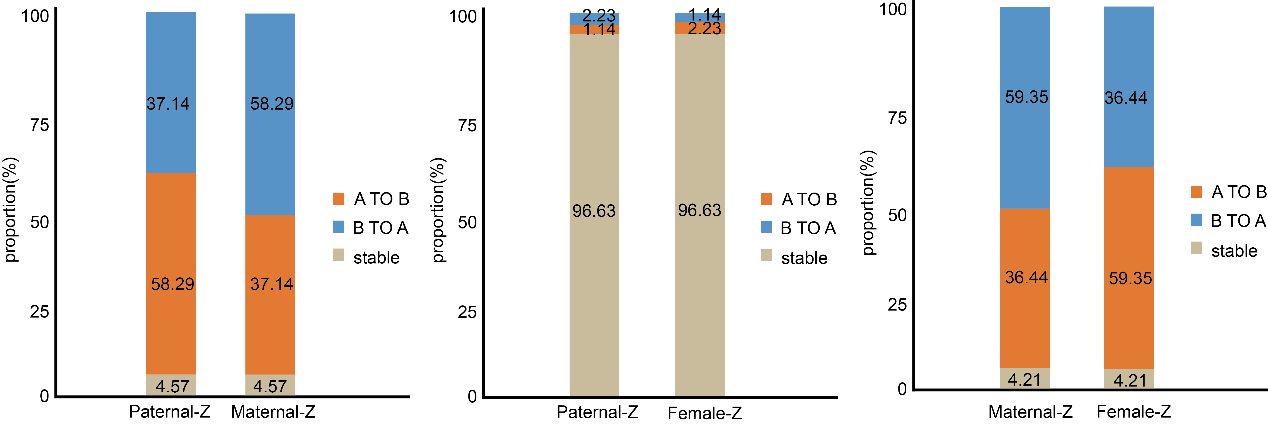

**S19 Fig. A/B compartment changes between paternal and maternal haplotypes.**

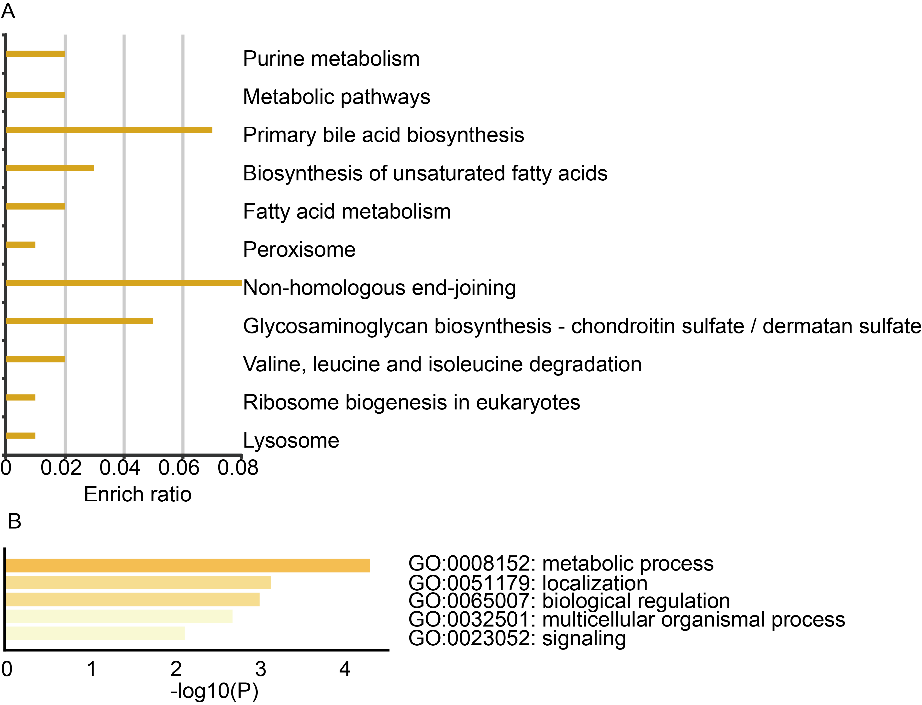

**S20 Fig. Changes in genes within A/B compartments between paternal and maternal alleles, annotated with KEGG and GO.** (A) KEGG analysis. (B) GO analysis.

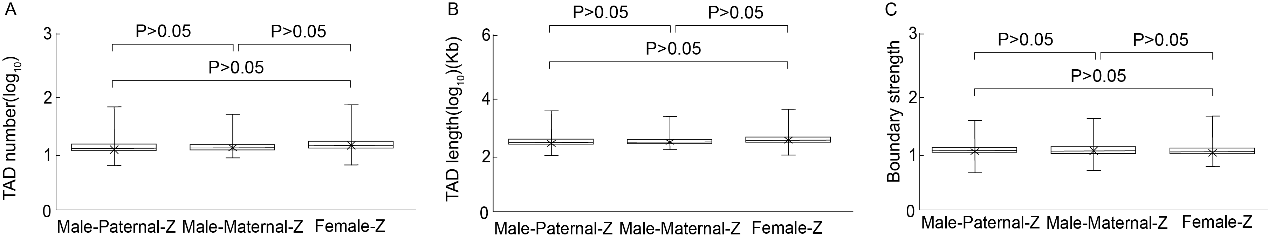

**S21 Fig. Statistical charts of haploid Z-chromosome TADs in Muscovy ducks.** (A) Comparison of TAD numbers on the haploid Z chromosome. (B) Length distribution of haploid Z-chromosome TADs. (C) Boundary strength comparison of haploid Z-chromosome TADs.

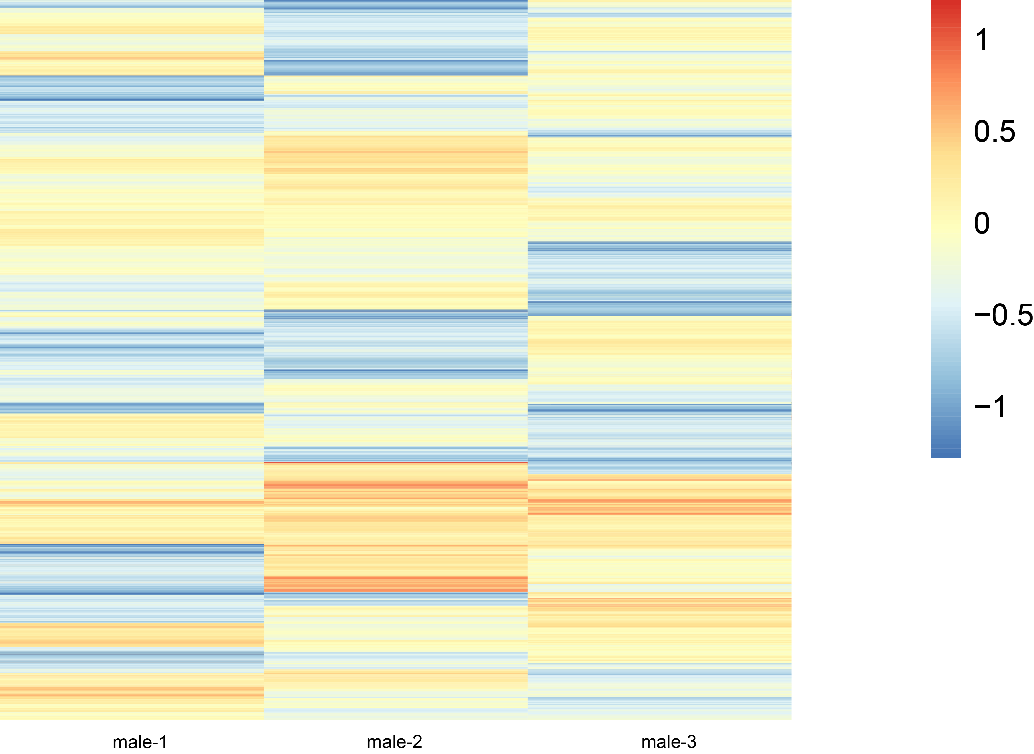

**S22 Fig. Haploid Z-chromosome ATAC-seq distribution map.** Differential heatmap of ATAC-seq peaks between paternal and maternal alleles in male individuals.

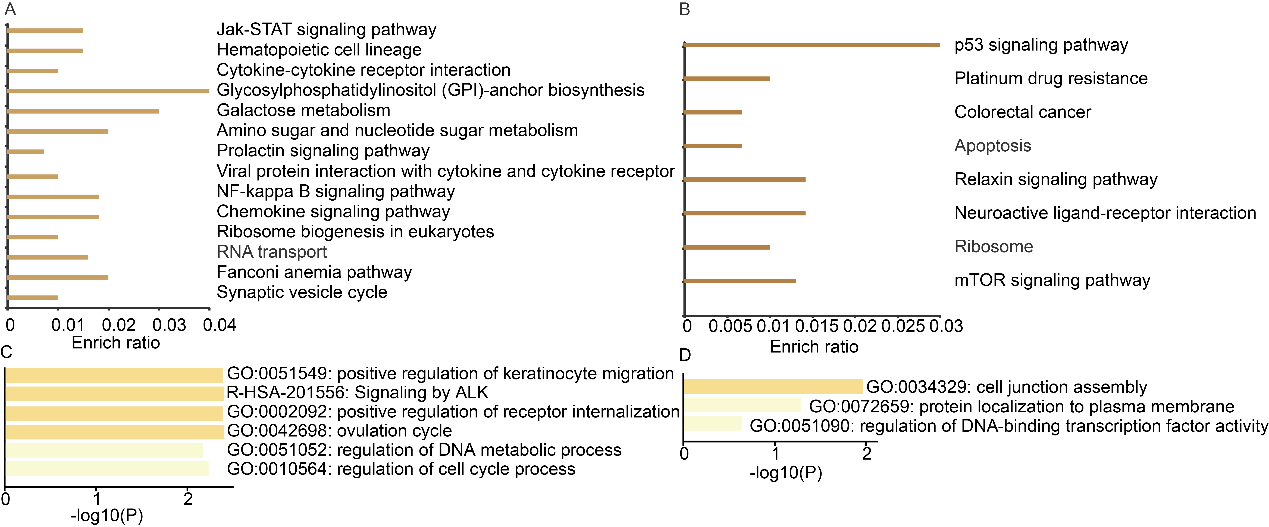

**S23 Fig.** **Identification of genes with allelically biased TSS peak clustering.** This analysis ultimately identified 40 genes with allelically biased TSS peak clustering, including two TSSs (*LOC112531540* and *MFSD5*) displaying exclusive paternal peak clustering on the Z chromosome and two TSSs (*LOC101750948* and *SCAMP1*) demonstrating maternal-specific peak clustering.

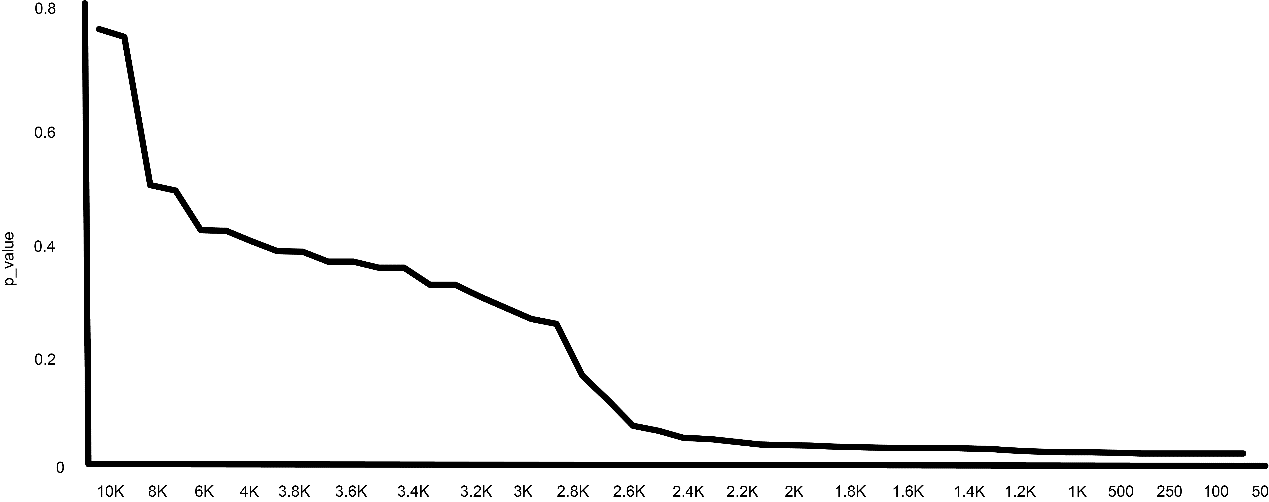

**S24 Fig. Sliding window segmentation of the whole genome.**

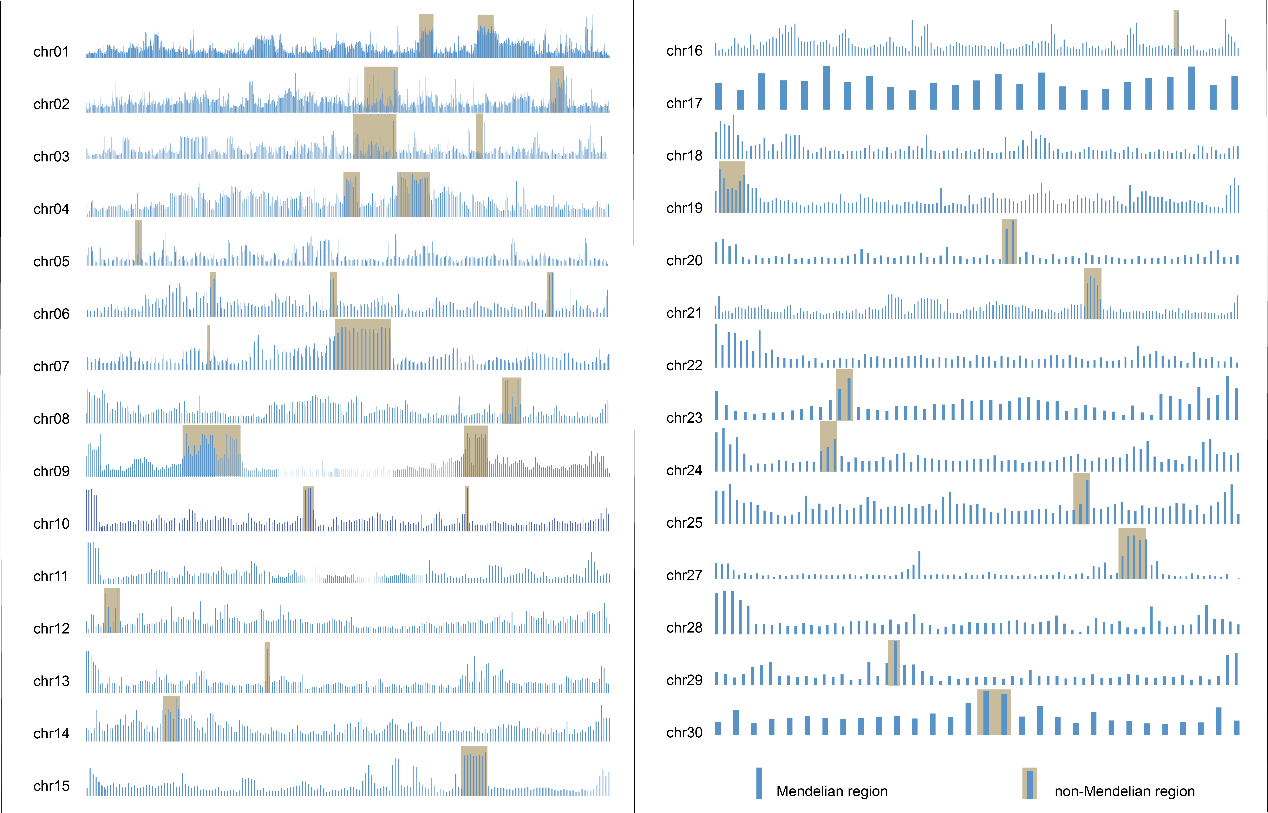
**S25 Fig. Genotype distribution map of the whole genome.** Genotypes were categorized into Mendelian and non-Mendelian regions based on the analysis of six haploid individuals and their parents.

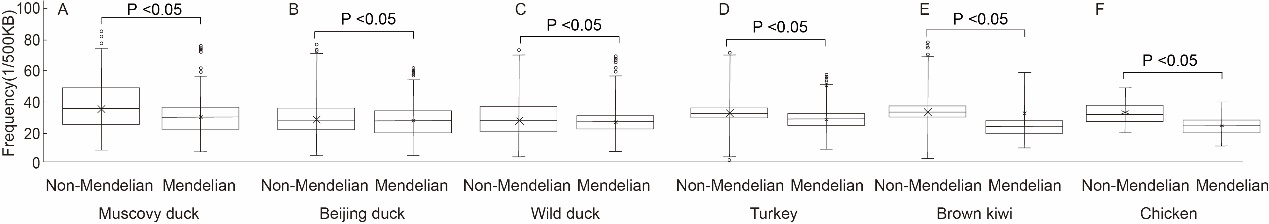

**S26 Fig**. **Non-Mendelian regions in different species.** Characteristics of non-Mendelian regions across different species. (A-E) Scatter plots showing the frequency of repeated sequences in Muscovy duck compared to Pekin duck, Mallard, Turkey, and Kiwi.

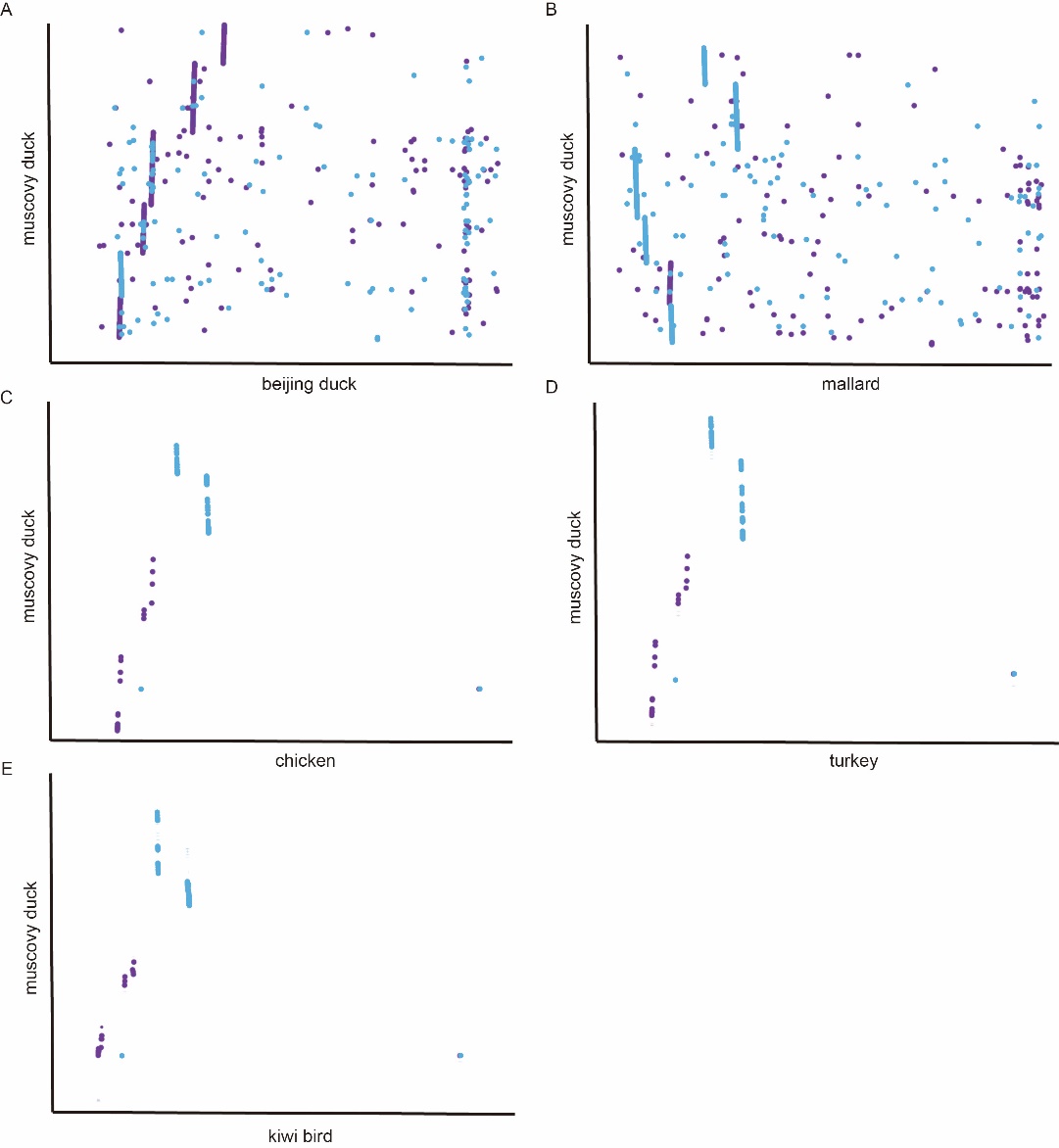

**S27 Fig. Syntenic analysis of motif enrichment.** Syntenic analyses demonstrate that the above four types of DNA motifs were also enriched in homologous regions of non-Mendelian regions from five divergent avian genomes (Chicken, Turkey, Mallard, Pekin duck, and Kiwi).
